# Chromosome-aware phylogenomics of Assassin Bugs (Hemiptera: Reduvioidea) elucidates ancient gene conflict

**DOI:** 10.1101/2023.03.22.533714

**Authors:** Alexander Knyshov, Eric R. L. Gordon, Paul K. Masonick, Stephanie Castillo, Dimitri Forero, Rochelle Hoey-Chamberlain, Wei Song Hwang, Kevin P. Johnson, Alan R. Lemmon, Emily Moriarty Lemmon, Samantha Standring, Junxia Zhang, Christiane Weirauch

## Abstract

Though the phylogenetic signal of loci on sex chromosomes can differ from those on autosomes, chromosomal-level genome assemblies for non-vertebrates are still relatively scarce and conservation of chromosomal gene content across deep phylogenetic scales has therefore remained largely unexplored. We here assemble a uniquely large and diverse set of samples (17 Anchored Hybrid Enrichment [AHE], 24 RNA-Seq, and 70 whole-genome sequencing [WGS] samples of variable depth) for the medically important assassin bugs (Reduvioidea). We assess the performance of genes based on multiple features (e.g., nucleotide vs. amino acid, nuclear vs. mitochondrial, and autosomal vs. X chromosomal) and employ different methods (concatenation and coalescence analyses) to reconstruct the unresolved phylogeny of this diverse (∼7,000 spp.) and old (>180 MYA) group. Our results show that genes on the X chromosome are more likely to have discordant phylogenies than those on autosomes. We find that the X chromosome conflict is driven by high gene substitution rates that impact accuracy of phylogenetic inference. However, gene tree clustering showed strong conflict even after discounting variable third codon positions. Alternative topologies were not particularly enriched for sex chromosome loci, but spread across the genome. We conclude that binning genes to autosomal or sex chromosomes may result in a more accurate picture of the complex evolutionary history of a clade.

## Introduction

Advances in DNA sequencing and bioinformatics have made analyses of large-scale phylogenetic matrices much more feasible. While these large datasets have resolved some relationships among major animal lineages (Dunn et al. 2008), many others remain contentious (Rodríguez-Ezpeleta et al. 2007; Smith et al. 2015; Murphy et al. 2021). The large size of phylogenetic matrices mainly limits stochastic errors, which were pervasive in the small-scale Sanger-based datasets of the past (Young and Gillung 2020). In contrast, other types of errors, known as systematic or non-random bias (Kapli et al. 2021), appear to remain unaffected by an increase in the amount of sequence data analyzed (Jeffroy et al. 2006; Rodríguez-Ezpeleta et al. 2007; Philippe et al. 2011; Kumar et al. 2012). This realization has promoted the development of new approaches to disentangle conflicting relationships (Arcila et al. 2017; Simon 2020).

Systematic bias encompasses several methodological factors (Simion et al. 2020) and biological phenomena. Among the latter, incomplete lineage sorting (ILS) and hybridization are commonly discussed as underlying causes of gene conflict (Rannala et al. 2020). Another source of phylogenetic conflict, chromosomal linkage, has so far received much less attention (Fontaine et al. 2015; Li et al. 2019). Recent advances in genome sequencing now allow for the linkage of largely complete but discontinuous genome assemblies into chromosomal scaffolds (Dudchenko et al. 2017; Yamaguchi et al. 2021). Probing the mixed phylogenetic signal across genomes in phylogenomic analyses has uncovered that gene features such as chromosomal linkage and GC content, may be predictive of phylogenetic signal, at least in certain mammal lineages (Li et al. 2019).

Assessing the chromosomal linkage of phylogenetic markers can potentially help explain observed systematic gene conflict. Sex chromosome genes often differ from autosomal genes in their evolutionary rate (Wilson and Makova 2009a; Wilson and Makova 2009b). A common pattern termed the “faster X effect”, is an elevated substitution rate of coding loci on the sex chromosome in the homozygous sex (i.e., X or Z; also called “hemizygous” sex chromosomes for brevity, herein) compared to autosomes (Mank et al. 2007; Meisel and Connallon 2013; Oyler-McCance et al. 2015). Hypotheses explaining this phenomenon focus on disparities in selection, population size, and expression between the sex chromosomes and autosomes (Meisel and Connallon 2013). In contrast, other factors, such as lower rates of mitotic division and recombination, may promote a relatively slower rate of change for X-linked loci. In cases of extensive ancient hybridization, a lower rate of recombination can enrich portions of the homozygous sex chromosome for loci that more accurately reflect the speciation history of a clade (Li et al. 2019). The relative importance of these conflicting pressures differs across taxa (Xu et al. 2012) but in most vertebrates studied, the sex chromosome in the homozygous sex appears to have a faster evolutionary rate than autosomes. Thus, loci located on the sex chromosome may be better suited for estimating shallower divergences than loci found on autosomes.

Although the X chromosome was observed for the first time in the true bug *Pyrrhocoris apterus* in 1891 (Paliulis et al. 2023), the genomic study of X chromosomes in invertebrates has lagged behind vertebrates. Relatively few chromosomal-level genome assemblies are available to assess the prevalence of a faster X effect across taxa, and the conservation of chromosomal gene content across deep phylogenetic scales is largely unknown. Among the existing studies on invertebrates, some (e.g., those on moths, spiders, and Sternorrhyncha) have shown the hemizygous sex chromosome with symptoms of the faster X effect (Sackton et al. 2014; Bechsgaard et al. 2019; Li et al. 2019), while in others the hemizygous chromosome evolves at the same or slower rate than the autosomes [beetles (Whittle et al. 2020) and stick insects (Parker et al. 2022)]. Studies on *Drosophila* have been conflicting, with only some supporting a faster X effect, which is generally weak (Betancourt et al. 2002; Counterman et al. 2004; Thornton et al. 2006; Hu et al. 2013). In damselflies with rampant hybridization, the X chromosome is particularly resistant to introgression (Swaegers et al. 2022). Fewer studies have assessed the depth of chromosome gene content conservation in invertebrates. In Diptera, there are major differences in the number of X-linked genes across lineages, driven by the convergent evolution of paternal genome elimination (Anderson et al. 2022). However, other studies have observed a relatively high level of conservation of gene content extending across the ordinal level (Meisel et al. 2019; Li et al. 2020; Li et al. 2022), with up to 25% of genes shared across more than 400 million years divergence in some cases. Long term conservation of gene content is necessary to observe any concerted effect of evolution of loci linked to sex chromosomes at deep phylogenetic scales.

An ideal group to further the study of X chromosome related evolutionary processes is Reduvioidea, the assassin bugs and relatives. This group comprises one of the most speciose lineages of Heteroptera (Schuh and Weirauch 2020) and is distributed worldwide (Maldonado Capriles, 1990). Assassin bugs are predominately predatory and display a great diversity of morphological and behavioral specializations (Weirauch et al. 2014). The group comprises two families, the species-poor Pachynomidae (∼30 spp.; 2 subfamilies) and the diverse Reduviidae (∼7,000 spp.; 24 subfamilies), with the latter including the blood-feeding and medically relevant kissing bugs (Triatominae), the vectors of Chagas disease. Despite being one of the largest superfamilies of true bugs and drawing the attention of medical entomologists, the evolutionary history of Reduvioidea is understudied. Phylogenetic hypotheses across this lineage include a morphological study (Weirauch 2008), Sanger-sequencing based analyses (Weirauch and Munro 2009; Hwang and Weirauch 2012), as well as a phylogenomic study with limited taxon sampling (Zhang, Gordon, et al. 2016). Only the analysis of Hwang and Weirauch (2012) sampled Reduviidae relatively comprehensively and recovered many subfamilies with high support, but largely failed to resolve intersubfamilial relationships. In contrast, the sole data-rich (370 loci) analysis to date (Zhang, Gordon, et al. 2016) sampled only 14 of the 26 reduvioid subfamilies. This analysis detected phylogenetic conflict, manifested in differences between concatenation- and coalescence-based analyses, demanding further investigation. To date, phylogenetic conflict across this old group [>180 MYA; (Johnson et al. 2018)] has not been investigated, partly because genome-scale data for Reduvioidea have remained scarce and the only available chromosome-level annotated assemblies originate from two closely related species of Triatominae (Mesquita et al. 2015; Liu et al. 2019).

Here, we investigate the phylogenetic conflict among loci through an assessment of their sex chromosomal linkages while reconstructing the evolutionary history of Reduvioidea based on a large phylogenomic dataset (2,286 loci; 23 of the 26 subfamilies). We sequenced 84 species of Reduviidae using hybrid capture (Lemmon et al. 2012) and genome skimming (Zhang et al. 2019) approaches. These two data types were combined with existing RNA-Seq and reference genomes in a single streamlined pipeline. Relying on available chromosome-level genomic assemblies, we interpolated linkage of loci in other taxa and compared the phylogenetic signal of the X chromosome with that of autosomes. Going beyond auto-sex discordance, we investigated and attempted to interpret results in the context of recent advances in gene conflict interrogation by conducting nucleotide- and amino acid-based analyses, comparing concatenation and coalescence-based analyses, and discerning clusters of genes with common phylogenetic signal.

## Results

### Dataset construction

To assemble the phylogenetic dataset, we combined sequences derived from Anchored Hybrid Enrichment (AHE, Lemmon et al. 2012), RNA-seq, and low-coverage whole genome sequencing (WGS) approaches, as well as high quality reference genomes, using an in-house developed bioinformatic pipeline (Fig. 1). We mined protein-coding AHE loci from all data types, resulting in a sub-dataset (“AHE dataset”) with 111 taxa (Table S1), 381 loci, and 231,153 positions in the nucleotide (NT) matrix (77,051 positions in the amino acid (AA) matrix). For 94 samples with transcriptomic and genomic data we obtained additional separate orthologous loci using the results of an OrthoMCL analysis (Gordon 2017). After filtering OrthoMCL-obtained orthogroups based on a number of criteria, this additional dataset (“OMCL dataset”) had 1,905 loci, and 1,566,147 nucleotide positions in the NT matrix (522,049 AA positions in the AA matrix). We used the combined AHE+OMCL dataset (111 taxa, 2,286 loci, and 1,797,300 nucleotide positions; 599,100 AA positions) for phylogenetic analyses. We split loci based on their linkage (X vs. autosome) in the triatomine reference taxa (Liu et al. 2019). The autosomal (AU) dataset comprised 2,141 loci and 1,690,446 NT positions and the X chromosome subset 145 loci and 106,854 NT positions. We further confirmed X-linkage of loci across reduviids as described in the “Chromosomal linkage of loci” section below. To investigate possible discordance between nuclear and mitochondrial phylogenetic signals, we attempted to obtain mitochondrial genomes (partial or complete) from all data types. Because of missing data distributions, only protein-coding mitochondrial genes were selected for downstream analysis, resulting in a mitochondrial dataset (“MT dataset”) containing 101 taxa and 16,008 nucleotide positions. For a separate analysis of entire gene content, we analyzed a full set of orthogroups that were detected by OrthoMCL using presence/absence coding (Pett et al. 2019). Despite some methodological shortcomings of this analysis (in incomplete genomic assemblies “absence” could mean both gene loss and missing data), it constituted a useful yet computationally efficient approach to utilize the additional available genomic data in an attempt to find phylogenetic signal and X chromosome associated patterns in gene gain and loss events. The extended methodology and results of this expanded analysis are available in the supplements (Supplemental Text S1).

**Figure 1.**
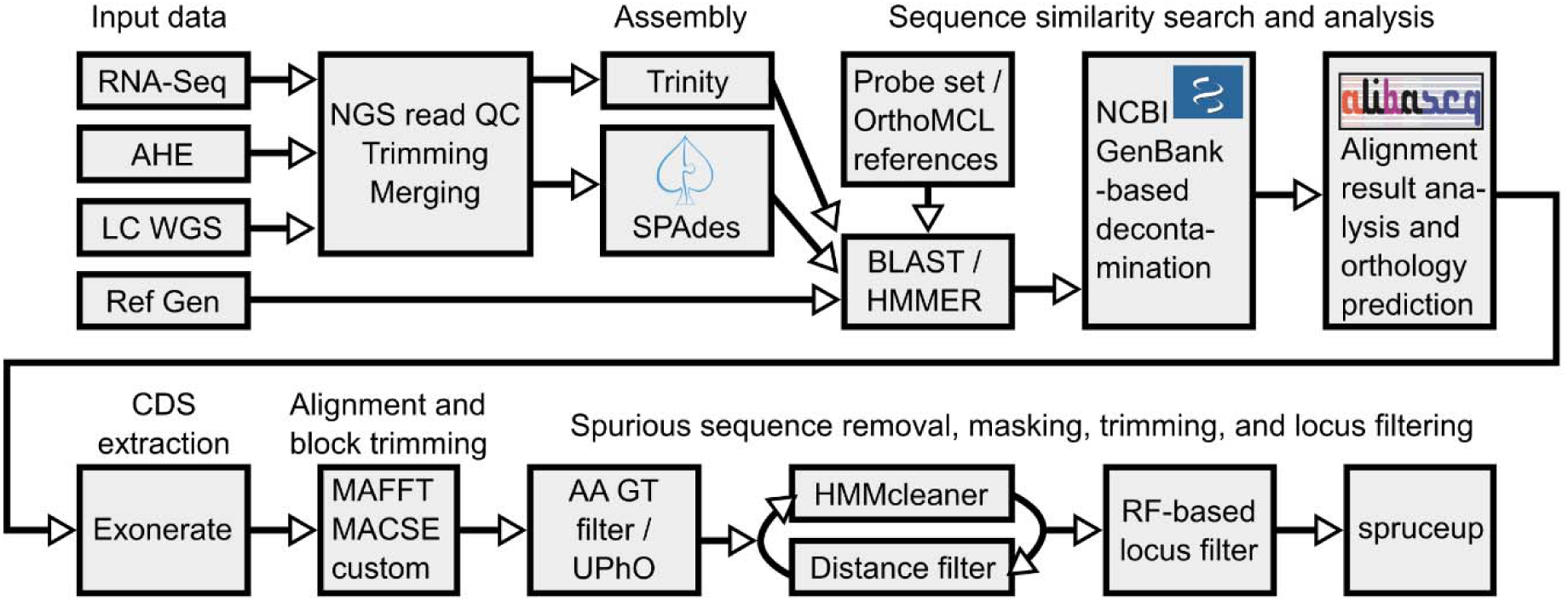
Schematic of the pipeline to produce the set of loci used in the study. Raw reads were run through quality control and preprocessing, followed by a de novo assembly. Obtained assemblies, together with three reference genome assemblies, were searched for homologs of the AHE and OMCL loci and extracted using *ALiBaSeq*. *Exonerate* was then used to precisely extract CDS, which were aligned and block-trimmed on protein level. Amino acid-based gene trees were used to remove distant outliers, and in case of the OMCL dataset, more precisely detect orthologs. Two rounds of segment trimming and further sequence outlier removal were performed, followed by removing loci with extreme RF-distance to the species tree, and concluded by a final segment trimming of the entire matrix using *spruceup*.

### Phylogenetic relationships and conflict

Six primary phylogenomic analyses were conducted to reconstruct the evolutionary history of assassin bugs (Figs 2, S1). We first analyzed the AHE+OMCL dataset in concatenation, using either the NT (“NT analysis”) or AA matrix (“AA analysis”), as well as with a coalescence-based approach using nucleotide-based gene trees in Astral (“Ast analysis”). The MT dataset was analyzed separately in a concatenation-based framework (“MT analysis”). We then analyzed putatively autosomal (“AU analysis”) and X-linked loci (“X analysis”) separately to identify possible phylogenetic discordance between these datasets (Fig. 2). Our analyses unambiguously recovered monophyly of both Reduvioidea (N1; full support in all datasets) and Reduviidae (N2; full support except X analysis). Similarly, Higher Reduviidae (N4) were fully supported in all analyses while the Phymatine-complex (N3) was not recovered in the X analysis; these two clades represent the deepest split in the phylogeny of Reduviidae. A number of subfamilies were recovered as monophyletic across all (i.e., Hammacerinae, Phymatinae, Peiratinae, Vesciinae, Stenopodainae, and Bactrodinae), or all except the X and/or MT analyses (Holoptilinae, Ectrichodiinae, Triatominae, and Salyavatinae). Consistent with published hypotheses, the large subfamily Reduviinae (>1,100 described spp.) is highly polyphyletic with many taxa currently recognized as reduviines recovered as distantly related lineages. Also corroborating published phylogenies, Emesinae were consistently rendered paraphyletic by Saicinae and Visayanocorinae, and Harpactorinae by Bactrodinae. Few intersubfamilial relationships were recovered with full or high support across all or most analyses. Examples are Hammacerinae as sister taxon to the remaining Phymatine-complex assassin bugs (all except X analysis with full support) and relationships among Stenopodainae, Triatominae and the *Zelurus* group of Reduviinae (all except MT analysis with full support). With regard to novel hypotheses, “Harpactorinae” + Bactrodinae formed the sister taxon to the small, subcorticulous reduviine *Heteropinus mollis* in all analyses. Also, in all analyses except X and MT, this clade was recovered as sister taxon to the reduviine genera *Nalata* and *Microlestria*, Epiroderinae, and Phimophorinae. Finally, the large clade (N7) comprising Cetherinae, Chryxinae, Pseudocetherinae, Salyavatinae, and the bulk of genera classified as Reduviinae also generally received high support across our analyses.

**Figure 2.**
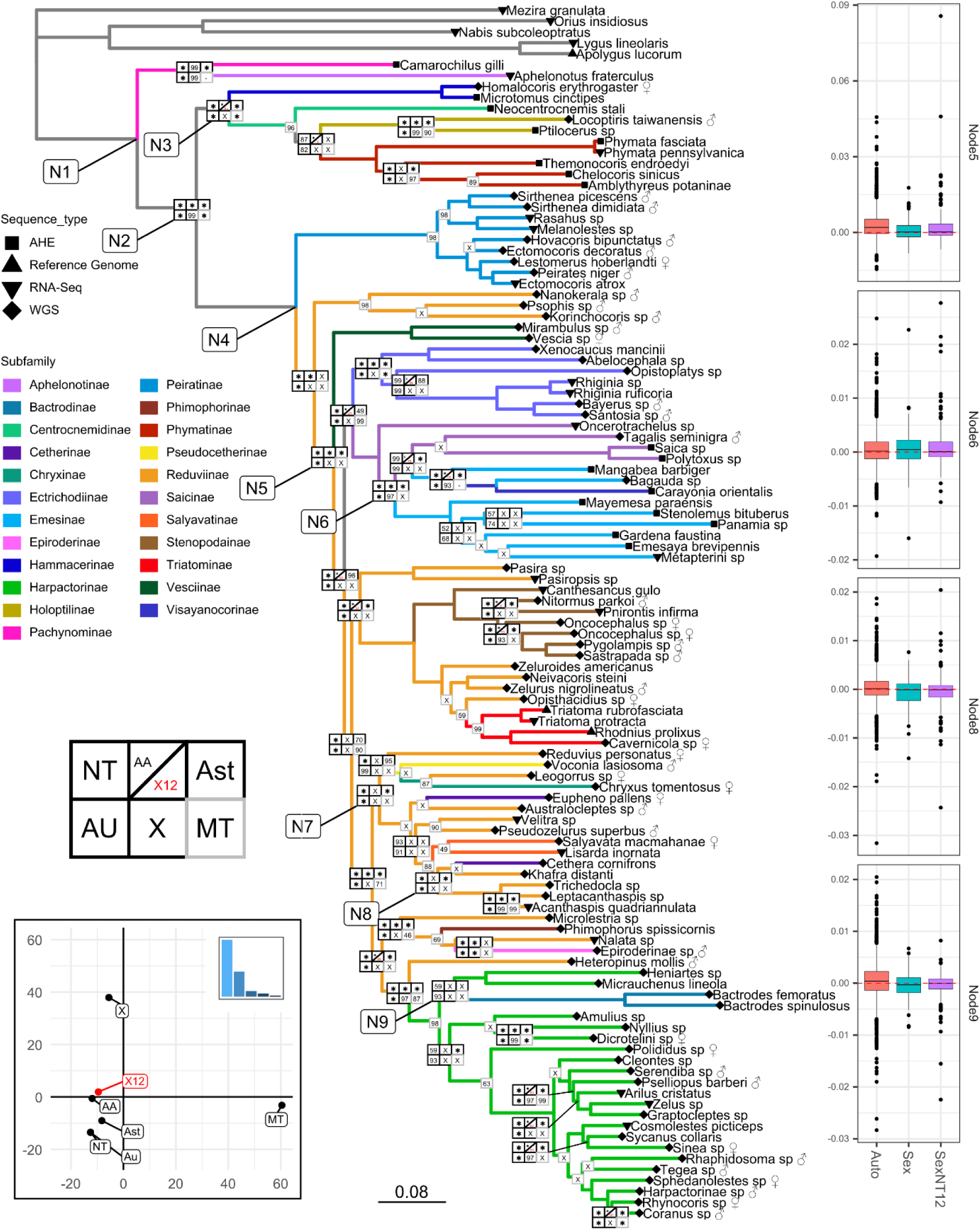
The topology in center was inferred based on the NT matrix of the combined AHE+OMCL dataset. Branches are colored based on the current subfamiliar classification of the 106 ingroup taxa. Tip symbols represent data type, sex is annotated for genomic taxa with morphological sex determination. Node boxes represent UFBoot (IQ-TREE-based analyses) or local posterior probability (Astral-based analyses) support values in different analyses (asterisk denotes full support in a given analysis) or include X when a node was not recovered relative to the NT analysis or by a dash “-” when a node could not be recovered due to reduced taxon sampling. Only nodes with a conflict or with less than 100% support in any analysis have nodal plots shown. When only the mitogenome dataset had less than 100% support, it was shown by itself in the interest of conserving space. From the distribution of the nodal boxes it is evident that the conflict between analyses primarily concerns the backbone of Reduvioidea and several deep divergences within subfamilies, while most recent divergences are supported across all analyses. Additionally, X chromosome and mitochondrial datasets are the two most discordant. Lower left panel represents an RF-based PCoA analysis of topologies obtained from different analyses, only first two PCs shown. Results of this analysis show that both X chromosome and mitochondrial topologies are drastically different from the rest, however also different from each other. Additionally, the X12 dataset (with third codon position removed) is relatively similar to full and autosomal analyses. Bar plots on right show likelihood difference between a given node shown in the phylogeny (N5-6, 8-9) and next most likely topology with a different relationship around that node for each gene (positive scores mean that the next most likely tree with an alternative node relationships is more likely for most genes). Results show that, for several of the nodes, correcting X signal (X12) made the likelihood difference distribution closer to that of autosomal genes. Additional nodes labeled (N1-4, N7) are discussed in text.

Despite many of the nodes receiving full support in the NT-based analysis, we observed considerable conflict between the NT topology and the topologies derived from the AA, Ast, X, and MT analyses. For instance, Peiratinae were the earliest diverging subfamily among the Higher Reduviidae in the NT analysis, but are sister either to the *Psophis* group of Reduviinae in the Astral and MT analysis, or to a part of Ectrichodiinae in the X analysis. While such conflict is ubiquitous along the generally less-well supported backbone of Reduviidae, some conflict was also observed within well-supported clades. For example, in the NT and AU analyses Bactrodinae were recovered as sister taxon to Apiomerini (represented by *Heniartes* and *Micrauchenus*), while AA, X, and MT showed Bactrodinae as the sister to the Higher Harpactorinae, or to Dicrotelini plus Higher Harpactorinae.

### Chromosomal linkage of loci

Among the analyses, the conflict between the loci putatively associated with the X chromosome (X analysis) and the autosomal loci (AU analysis) was particularly noticeable. To confirm that X-chromosomal loci of Triatominae are also X-linked in other reduviids, we investigated conservation of X-located loci in our taxon set. Li et al. (2020) and Mathers et al. (2020) showed that X linkage of loci was conserved between two closely related blood-feeding reduviids, *Triatoma rubrofasciata* and *Rhodnius prolixus*, as well as that sex chromosomal loci had stable linkage in other hemipteran groups. Since no other chromosome level assemblies of Reduviidae are available, we used recently published chromosomal assemblies of *Apolygus lucorum*, a distant outgroup of Reduvioidea, to verify sex chromosome loci conservation on a deeper evolutionary scale. We followed the methods of Li et al. (2020) to determine the conservation of synteny between *Apolygus* and *Triatoma*. Results of the analysis (Fig. 3a) suggested that blocks of genes from the X chromosome of *Triatoma rubrofasciata* reciprocally matched to chromosome 1 (presumably the X chromosome) of *Apolygus lucorum*, with no sex chromosomal blocks matching to any autosomes. This is the first indication that X chromosome loci might be conserved at least to some extent in their chromosomal linkage across Cimicomorpha [∼225 Mya (Johnson et al. 2018)], a much deeper level than what has been previously shown [22, 32, and 57 Mya in different hemipteran lineages (Mathers et al. 2020)].

**Figure 3.**
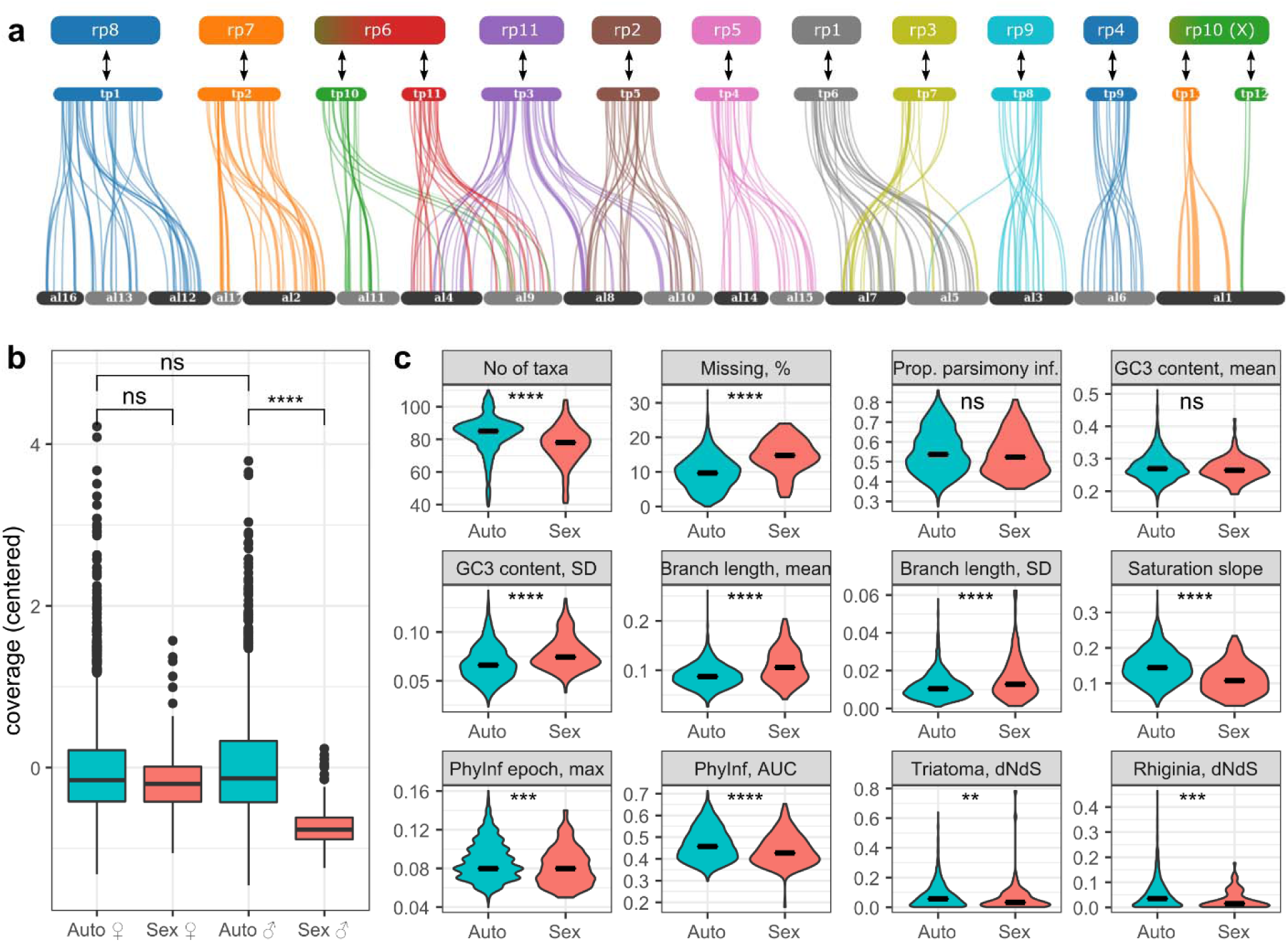
The top panel (a) shows the results of the gene synteny analysis between *Triatoma rubrofasciata* (tp) and *Apolygus lucorum* (al) chromosomes, with *Rhodnius prolixus* chromosomal synteny (rp) shown at the top. The analysis shows fairly conserved synteny between groups of al chromosomes and tp chromosomes. More interestingly, no syntenic blocks for X loci of tp were found on autosomes of al. Bottom left panel (b) shows the distribution of average locus depth in the combined AHE+OMCL dataset on autosomes and X chromosome by morphological sex determined for taxa with sex symbols indicated on Fig. 2. Results show male X chromosomal loci having significantly lower average coverage than female X loci or male autosomal loci. Bottom right panel (c) shows the distribution of various locus properties in autosomal and X chromosome loci, outliers were excluded, for the full data see Fig. S2. Most properties have significantly different distributions, with the exception of proportion of parsimony informative sites and mean GC content of the third codon position.

To confirm that the 145 putatively X-located loci share the same chromosome in other ingroup taxa, we determined locus coverage depth of all loci, and compared it between specimens with identified biological sex. Results showed (Fig. 3b) that putatively X-located loci had significantly lower coverage in males compared to females. X-located loci in males also had lower coverage compared to autosomal loci in both sexes. Based on these results, we proceeded to treat the 145 putatively X-associated loci as sex chromosome located loci, with all other loci being treated as autosomal. Since the X chromosome was the only one with conserved gene content and thus with inferrable loci, and for simplicity reasons, we refer to classification of loci as autosomal or X-linked as “chromosomal linkage” in this paper.

### Autosomal and sex-chromosomal loci: phylogenetic signal and differences in properties

Using identified autosomal and sex chromosome loci, we performed phylogenetic reconstructions as well as a gene-by-gene log-likelihood fit difference analysis (Lee and Hugall 2003; Shen et al. 2017) while binning loci according to their chromosomal linkage. We rescaled log-likelihood differences by gene length to obtain per-base phylogenetic signal metric and avoid disproportional influence of long loci. Phylogenetic results showed (Fig. 2) considerable topological differences between autosomal and sex chromosomal topology, some of which we discussed above. Additionally, the X-chromosomal tree had a larger total tree length, suggesting an on average higher substitution rate of sex loci (Fig. S1). Results of the log-likelihood fit difference analysis showed that for several conflicting relationships (Fig. 2, N5-6, N8-9) there was a difference between median autosomal and X likelihood difference between tested topological rearrangements. Consistently with phylogenetic inference results, median X chromosome likelihood fit difference was either positive but lower than autosomal (favoring same topology but less strongly, less informative compared to autosomal dataset, N5); or positive but higher than autosomal (favoring same topology more strongly, potentially could have more decisive impact on combined topology inference, N6); or negative (favoring opposite topology from the autosomal loci, N8-9).

We scored several locus properties in both autosomal and X-located loci to determine if any of the properties differ and could explain the observed gene conflict (Fig. 3c). Besides a larger proportion of missing data, which could be associated with lower coverage of X-loci in males, sex loci also had higher substitution rates and levels of saturation. However, selection analyses showed no significant difference in dn/ds ratio between sex chromosome and autosomal genes. All loci selected for phylogenetic inference appear to be under relatively strong purifying selection. Mean GC content of third codon positions (GC3) was similar between X-located and autosomal loci, however GC3 variation of X was significantly higher.

### Correcting the misleading sex chromosomal signal

Since assessment of both phylogenetic inference and locus properties pointed to possible saturation resulting from high substitution rate in the presence of stabilizing selection, we hypothesized that the observed phylogenetic signal of X-located loci is artefactual. In an attempt to correct this signal, we excluded 3rd codon positions, which are the most impacted by saturation, and reanalyzed the datasets. The analysis of X loci with the exclusion of 3rd codon position (X12) yielded a topology much more congruent with that of autosomal loci and very similar to the AA-based tree (Fig. 2, PCoA plot, Fig. S1). We also reanalyzed likelihood fit difference for the loci in the corrected dataset. Results showed (Fig. 2, bar plots), that the corrected sex-chromosomal signal was much more in line with the autosomal signal in the four previously contentious nodes (i.e, median log-likelihood fit difference values between conflicting topologies became more similar between autosomal and X12 assessments).

### Gene conflict beyond the sex chromosome

Much of the strong gene conflict was alleviated by addressing saturation artifacts of the X-located loci (Fig. S1 XNT12; e.g., position of Peiratinae), but several relationships remained ambiguous (e.g., position of Bactrodinae). To investigate gene conflict beyond chromosomal linkage, we computed gene tree distances and ran a PCoA analysis of the distances. Seven clusters of genes (or ‘groves’) based on their topological similarities were identified (Fig. 4). On average, clusters contained the same proportion of sex genes as the entire dataset (6%). However, the most closely clustered group, grove 6, had fewer X genes (3.3%) while another cluster, grove 3 was uniquely enriched for X genes (13.5%). One of the clusters contained only three genes and was excluded from subsequent investigations. For each of the remaining clusters we inferred an ML tree, concatenating corresponding loci (N=86–766). Resulting topologies showed (Fig. 4; groves 1–6) that each cluster tree had a uniquely different position of one or several contentious taxa or clades in addition to contentious nodes shared between several groves (e.g., position of Bactrodinae, relationships within Emesinae and between Ectrichodiinae and Emesinae, and relationships within clade N7).

**Figure 4.**
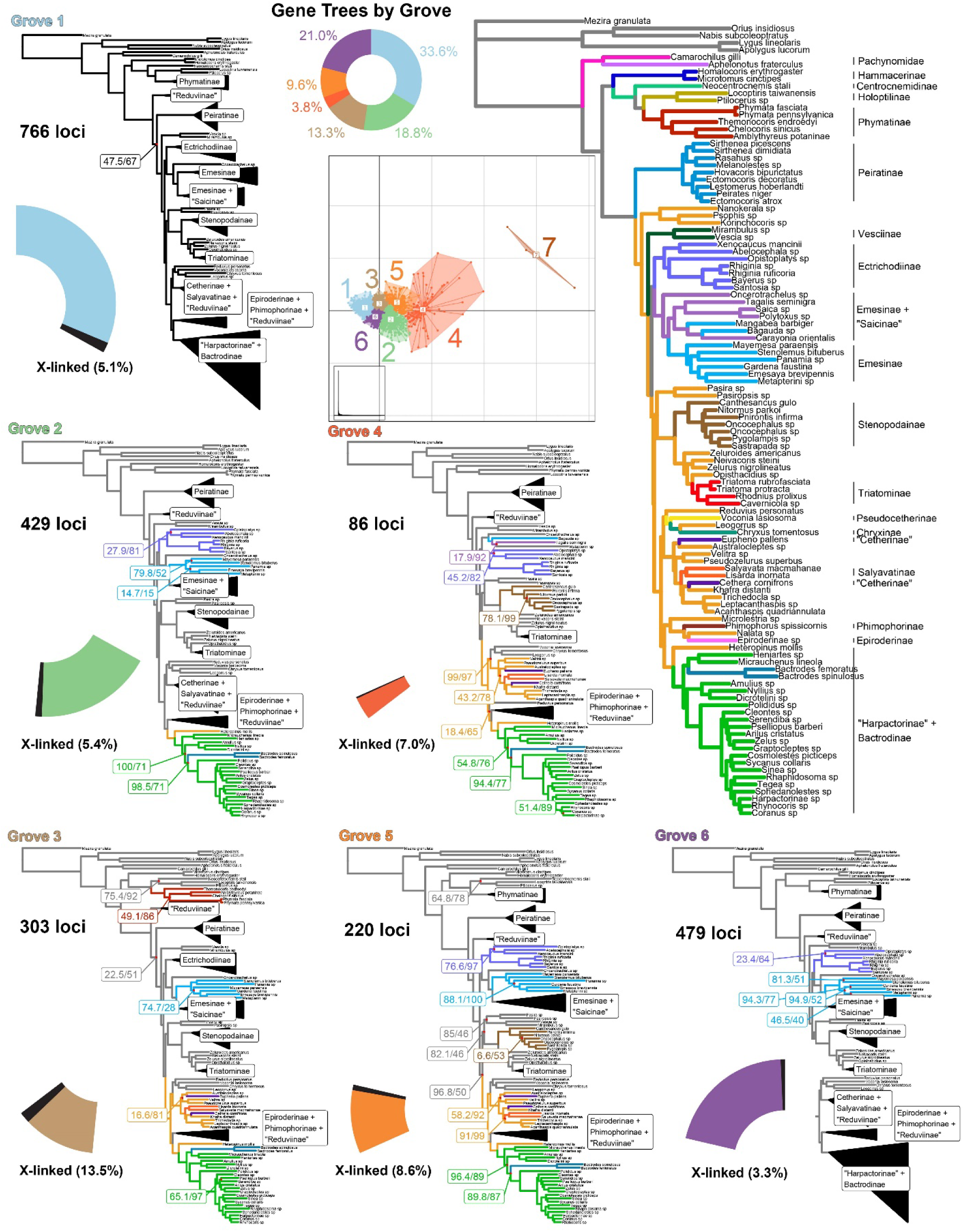
Visualization of gene tree conflict. NT12 gene trees were clustered into groves based on weighted RF distance calculated across gene tree phylogenies with nodes supported by less than 50% bootstrap collapsed using TreeSpace (PCoA). Analyses of full NT123 grove sets with IQ-TREE are shown; only nodes which differ from the combined Autosomal NT123 tree (large tree on right) include support values and relevant clades which contain differences are highlighted in color for emphasis. The number of loci in each grove is represented as a portion of a pie chart with the number of X-linked loci highlighted in black.

## Discussion

### Chromosomal linkage of loci can explain part of the conflict among loci

In a large phylogenomic analysis of assassin bugs and relatives (Heteroptera: Reduvioidea), we detected substantial phylogenetic conflict among nuclear loci. This conflict can at least in part be explained by loci associated with the X chromosome (Fig. 2). Despite only about 6% of the dataset being composed of genes associated with the X chromosome, several nodes in the combined analyses received lower support than in autosome-only analyses because of the auto-X signal conflict. Moreover, the X-only tree had a drastically different topology and branch length distribution from other subsets of data, driven in part by a higher substitution rate of X-linked loci. The matrix of X-linked loci also had more missing data, partially due to lower coverage of X-linked loci in males but also possibly impacted by higher genetic distance from the reference. Although larger distances probably did not impact homology searches, they can cause more trimming and masking during filtering steps in the pipeline. The fast substitution rate, however, could not be explained by positive selection, because there was no difference in dn/ds between autosomal and sex chromosomal partitions (Fig. 3c). However, sex loci were more saturated compared to autosomal markers, thus allowing us to hypothesize that it is the high substitution rate coupled with strong purifying selection that causes the discordance between X loc and autosomal loci trees. Thus, the X-loci discordant signal in our data is non-phylogenetic. The topology of X12 is highly congruent with the autosomal topology save for a few recalcitrant nodes. This congruence indicates that most of the conflict comes from homoplasy caused by highly saturated rapidly evolving genes and not from any shared strong conflicting signal retained by genes housed on the X chromosome with the same evolutionary history.

### Observed gene conflict is not solely explainable by X-loci signal

In most cases, phylogenetic conflict can rarely be attributed to one cause. In the case of phylogenetic conflict in Reduvioidea, X-chromosomal linkage and the associated high substitution rate only partially explains the observed discordance among loci. For example, some relationships with conflicting resolutions were retained in both full X and NT12 X analyses. X-linked loci were enriched in the genes comprising grove 3 (Fig. 4) and these together with autosomal genes in the grove could represent a group of loci with signal of hybridization or incomplete lineage sorting across both the autosomal and sex chromosome backgrounds. The mitogenome, similar to sex chromosomes, has a biased pattern of sex-based inheritance and we were curious to see if there was any group of loci that has been inherited in a similar way or share a similar signal. We did not observe any set of nuclear loci with the same phylogenetic signal as the mitogenome, which even with relatively limited signal, had strong conflicting resolution to all other examined trees. Alternative resolutions seen in other loci grouped into groves likely reflect true conflict. As chromosomal-level genomic assemblies become more prevalent, it may be possible to determine linkage of similar subsets of loci that would reflect explanatory introgression or hybridization events.

### Phylogeny of Reduvioidea

Our study presents the most data-rich and densely sampled phylogenetic hypotheses for the assassin bugs and relatives to date (111 taxa; 2,286 loci; 23 of the 26 reduvioid subfamilies). Of the three subfamilies missing from our analyses, one is thought to be part of the Phymatine-complex based on morphological data [Elasmodeminae (Carayon et al. 1958)], another rendered Salyavatinae paraphyletic in Sanger-based analyses [Sphaeridopinae (Gordon and Weirauch 2016)] and the last, Manangocorinae are monotypic, only known from the holotype, and morphology suggests that this subfamily may belong to clade N7. Topologies recovered from different datasets in our analyses are similar to but not identical with published hypotheses that attempted to resolve relationships across reduviid subfamilies (Weirauch and Munro, 2009; Hwang and Weirauch, 2012; Zhang et al. 2016) and to topologies from studies that aimed to uncovering relationships within subfamilies or clades of related subfamilies [Harpactorinae/Bactrodinae (Zhang, Weirauch, et al. 2016); Phymatine-complex (Masonick et al. 2017); Triatominae and relatives (Justi et al. 2014; Kieran et al. 2021); Ectrichodiinae (Forthman and Weirauch 2016)]. While several of the nodes along the backbone of Higher Reduviidae (N4) are unstable across our analyses, many aspects of our hypotheses are well-supported in the NT, AA, Ast, AU, and X12 analyses and can form the basis for a revised subfamily- and tribal-level classification built on diagnosable monophyletic groups. The polyphyly of Reduviinae, first documented by Weirauch (2008) based on morphology and corroborated by Sanger-derived (Hwang and Weirauch, 2012) and phylogenomic datasets (Zhang et. al., 2016) is exacerbated by recovering *Heteropinus* as a separate lineage and sister to the largely diurnal and vegetation-dwelling “Harpactorinae” + Bactrodinae. An assassin bug classification based on diagnosable clades will require the recognition of several of these distantly related reduviine lineages as separate subfamilies. In contrast, the paraphyly of “Harpactorinae” with respect to Bactrodinae, paraphyly of “Emesinae” and “Saicinae” relative to Visayanocorinae, and the relationships of reduviine taxa with non-reduviine subfamilies contained in clade N7 will require the synonymization of several currently recognized subfamilies. Combined phylogenomic and morphological datasets and analyses have been generated to formalize these changes to the classification (Masonick et al., in prep.; Standring et al., in prep.).

In addition to improving our hypothesis of the evolutionary history of reduvioids, newly sampled taxa in our study could aid in a more robust reconstruction of genomic patterns in the evolution of blood-feeding in Triatominae. Historically, comparative genomic studies on blood-sucking true bugs have been restricted to few distant independently parasitic taxa, without detailed comparison to the closest non-blood feeding relatives. However, studying the closest non-blood feeding outgroups can help more accurately pinpoint the genes involved in the feeding habit transition. For example, the *Rhodnius prolixus* genome possesses several odorant binding protein (OBP) gene clusters that were reconstructed to be uniquely duplicated in Triatominae given the taxon sampling (Mesquita et al. 2015). However, our gene content analyses (Supplemental Text S1) show that among the discovered OBP clusters only one (RproOBP13) is truly specific to Triatominae, with other OBPs recovered in non-blood feeding reduviids. This protein along with another that is not exclusive to Triatominae (RproOBP6) had previously been identified as expressed in the antennae of male and female insects (Oliveira et al. 2018) and thus likely plays a role in host-sensing.

### Investigation of X chromosome evolution is applicable to phylogenomics

Chromosome-level assemblies continue to be scarce and impede progress on understanding of sex chromosome evolution and examination of gene conflict. To gain knowledge more rapidly, phylogenomic data like ours can be combined with chromosomal assemblies to assess conservation of chromosomal loci and shed light on subsequent phylogenetic analyses. Although the prevalence of a fast X effect is not known across all arthropods, loci linked to X chromosomes are common in phylogenomic studies of arthropods (6% in our dataset but a cursory analysis of some other commonly used public locus sets in arthropods yielded similar proportions of X-linked loci, ∼5-10%). Gene content of X chromosomes appears conserved in some cases up to ∼200 Mya divergence between arthropod taxa, making gene content inferable for many small-scale phylogenetic studies with only distant relatives with a chromosomal-level assembly. Despite comprising a small proportion of loci, X-linked genes appear to have a considerable influence at least in the case of our dataset in which the tree includes rapid radiations and several groups have long branches. X chromosome loci can also have other features which would impact accurate phylogenetic reconstruction [e.g., relaxed purifying selection in aphids (Li et al. 2020) or stick insects (Parker et al. 2022)].

## Conclusion

In summary, we developed a method to organize four different kinds of heterogeneous genomic data types, achieving extensive taxon sampling to infer a phylogeny of this medically important group of arthropods. We were able to confirm a faster rate of evolution of X-linked loci in this group, which impacted accurate phylogenetic reconstruction but did not entirely explain other underlying gene conflict. Building upon these findings, we established a robust phylogeny of the group, highlighting contentious nodes with substantial conflict that will be valuable for understanding the evolution of assassin bugs. Furthermore, this approach is applicable to other organisms with only distant relatives with chromosomal-level reference assemblies to generate a more comprehensive understanding of sex chromosome and clade-specific evolution.

## Material and Methods

### Anchored Hybrid Enrichment

The Anchored Hybrid Enrichment (AHE) dataset consisted of 17 samples (Table S1). DNA from the samples was extracted using a Qiagen DNeasy Blood and Tissue kit. Library preparation, hybrid enrichment, and sequencing followed standard protocols (Lemmon et al. 2012). A Hemiptera probe set (Dietrich et al. 2017) was used for hybrid enrichment. Samples were sequenced on an Illumina HiSeq 2500 platform. Reads were processed with *clumpify* (BBMap v38.86 package) and deduplicated, then trimmed with *Trimmomatic* v0.36 (Bolger et al. 2014), merged with *bbmerge* (BBMap v38.86 package), error-corrected and assembled with *SPAdes* v3.12.0 (Bankevich et al. 2012).

### RNA-seq

RNA-seq data for the present study was originally obtained as part of the (Zhang, Gordon, et al. 2016). Briefly, RNA was extracted from head and thorax, or full body of the specimens, and cDNA libraries were prepared and sequenced at the W.M. Keck Center (University of Illinois) and AITBiotech PTE LTD (Singapore) using an Illumina HiSeq platform and paired-end 100-bp chemistry. Reads were trimmed using Trimmomatic (Bolger et al. 2014) and assembled with Trinity (Haas et al. 2013) as part of (Gordon 2017). For four outgroup taxa original assemblies from (Zhang, Gordon, et al. 2016) were used.

### Whole Genome Sequencing

DNA from the samples was extracted using either Qiagen DNeasy Blood and Tissue kit or using a combination protocol of Qiagen Qiaquick and DNeasy kits (Knyshov et al. 2018). Five samples (Dicrotelini sp., *Heniartes sp.*, *Cleontes sp.*, *Amulius sp.*, and *Bactrodes femoratus*) were subjected to a library prep protocol as in (Knyshov et al. 2018) and sequenced on a HiSeq X lane. DNA extracts of the remaining samples were used to prepare libraries as in (Lemmon et al. 2012), and the samples were sequenced on several NovaSeq6000 S4 lanes. Reads processing and assembly methods were the same as for the AHE samples.

### Previously available chromosome-level assembles

We also included chromosome-level assemblies of the two available reduviids, *Rhodnius prolixus* (Mesquita et al. 2015) and *Triatoma rubrofasciata* (Liu et al. 2019), and the assembly of *Apolygus lucorum* (Liu et al. 2021) as an outgroup. For *Rhodnius prolixus*, we used the chromosomal assembly provided by DNAZoo (Dudchenko et al. 2017) and also carried the unmasked original assembly (Mesquita et al. 2015) and predicted proteins (RproC3.3) through the initial steps of the pipeline to assess its performance as well as to compare the completeness of these data with the chromosome-level assembly.

### AHE dataset construction

Heteroptera bait regions of the Paraneoptera AHE kit used for enrichment were reprocessed to remove overlapping regions and combine exons of the same genes using a taxon used for probe design [similar approach was employed by (Van Dam et al. 2020)]. Coding sequence of the resulting regions was determined and translated amino acid (AA) sequences for each locus were obtained. The *Trinity*-based assembly of the taxon that was used for probe design (*Arilus cristatus*) was used as the reference. A total of 397 gene regions, representing original 478 AHE bait regions, were selected for downstream analyses.

Discontinuous megablast (*dc-megablast*) from the *BLAST* package (Altschul et al. 1990; Camacho et al. 2009) was used to search for the bait sequences in the RNA-seq (24 samples), WGS (67 samples), hybrid capture (17 samples), and chromosomal-level assemblies (3 samples). Due to noticeable contamination in some WGS and AHE samples, the matched contigs were prescreened for non-arthropod sequences via *megablast* against the NCBI database. Subsequently, non-contaminant matched contigs were searched against the reference assembly of *Arilus cristatus* using *dc-megablast* for the reciprocal best hit check. *ALiBaSeq* (Knyshov et al. 2021) was used to parse the *BLAST* results, perform reciprocal best hit check, stitch contigs of RNA-seq and WGS samples belonging to the same bait, and compile the resulting locus by locus FASTA files. We set *ALiBaSeq* to recover both matched and internal unmatched sequence regions (mode -x b) to allow for a more accurate CDS sequence extraction in the next step.

*Exonerate* v2.2.0 (Slater and Birney 2005) was used together with protein references of *Arilus cristatus* to accurately excise exons of the recovered sequences. Obtained CDS sequences were aligned on protein level with *MAFFT* (Katoh and Standley 2013). Sequences were trimmed on AA level. *MACSE* v2.03 (Ranwez et al. 2011) was used to transfer protein alignment and trimming onto nucleotide (NT) sequences. Following trimming, gene trees were reconstructed based on protein sequences using *RAxML* v8.2.12 (Stamatakis 2014) with the LG+G model. Excessively long branches representing likely paralogs were removed using a custom R script. This and subsequent filtering R scripts were based on functions from *APE* (Paradis and Schliep 2019), *Phangorn (Schliep 2011)*, and *SeqinR* (Charif and Lobry 2007) R packages. The filtered alignments were then carried two times through *HmmCleaner* v0.180750 (Di Franco et al. 2019) filtering, followed by a custom-made sequence distance filter to remove outliers. Loci shorter than 50AA (150bp) were discarded. Then NT alignments were used to reconstruct gene trees using *RAxML* with the “GTRGAMMA” model, which then had nodes below 33% Rapid Bootstrap Support (RBS) collapsed. Obtained trees were used to detect and remove cross-contamination as well as to filter weighted Robinson-Foulds (RF) distance (Robinson and Foulds 1981) outlier loci. Obtained filtered alignments were filtered with *spruceup* (Borowiec 2019). Concatenated NT and AA analyses were conducted in *IQ-TREE* v1.7-beta9 (Nguyen et al. 2015). Custom scripts, available at https://github.com/AlexKnyshov/PLS, were used to calculate per locus difference in log-likelihood for each Nearest Neighbor Interchange (NNI) (Lee and Hugall 2003; Shen et al. 2017). Loci with too many outlier nodes and outlier clades in particular loci were screened for homology errors and some removed.

### OrthoMCL-based dataset construction

We started with 32,675 orthologous clusters produced by (Gordon 2017). Briefly, 20 reduviid transcriptomes from (Zhang, Gordon, et al. 2016) along with the *Rhodnius* coding sequence were used as the input. The longest ORF was predicted using *Transdecoder*, and resulting sequences supplied to *OrthoMCL* (Li et al. 2003).

We queried the clusters in which at least 10 species have sequences and all orthologs in a cluster are single copy. We then only retained clusters passing the minimum length threshold (>=100AA) and lacking extraordinary mean pairwise distance (as those likely represented erroneous clustering). AHE genes were then found via BLAST and removed to avoid duplication of the AHE dataset. This filtering resulted in 2,296 clusters. Proteins for each orthogroup were aligned, trimmed from flanks, and *HMMER* profiles were generated. All sample assemblies were translated into six reading frames and searched for homologs using the profiles obtained earlier. Due to computational difficulties with performing a *HMMER* search on chromosome-level scaffolds of the three chromosome-level assemblies, *TBLASTN* search was used instead for these samples. The most complete sample from the initial OrthoMCL set (Pasir = *Pasiropsis sp.*) was selected as reference taxon for the RBH check. *ALiBaSeq* (Knyshov et al. 2021) was used to parse search results parsing, perform RBH check and sequence extraction.

Subsequent sequence processing was largely similar to AHE dataset. In order to mitigate elevated risks of paralog incorporation, in this dataset *ALiBaSeq* was used to pull out closely matching suboptimal hits in addition to the best hit, protein-based gene trees were reconstructed, and *UPhO* (Ballesteros and Hormiga 2016) was used to infer final orthologs.

### Mitochondrial dataset

Using available reduvioid references from GenBank, mitochondrial contigs were searched for in the whole read assemblies from above using dc-megablast. Recovered partial sequences were improved upon using a combination of *NovoPlasty* (Dierckxsens et al. 2017), *Mitobim* (Hahn et al. 2013), read mapping using *BBMap*, and final curation and annotation in *Geneious* (Kearse et al. 2012).

### Assessment of locus properties

We used *AMAS* (Borowiec 2016) and a modified gene_stats.R (Borowiec et al. 2015) to assess general locus stats such as length, number of taxa, proportion of parsimony informative sites. *RAxML* trees, computed for summary coalescence analyses, were used to assess average BS, average branch length, and saturation. Custom R scripts and APE package were used to compute GC content mean and variance across taxa per each codon position.

Final *IQ-TREE* analyses were used to record per site rates of all genes. These rates were used with the site_summer function of *PhyInformR* package (Dornburg et al. 2016) to reconstruct a phylogenetic informativeness curve (Townsend 2007). We sampled informativeness at 0.01 intervals as in the web application *PhyDesign (López-Giráldez and Townsend 2011)* and contrary to the default behavior of *PhyInformR* to only sample at nodes, as the latter led to aberrations in curve smoothing for some loci. We then recorded the epoch (normalized interval along tree height) of the max informativeness, as well as computed total informativeness as area under curve.

In order to evaluate the selection strength on the loci, we used three pairs of closely related species across the reduvioid tree: *Phymata pennsylvanica* and *Phymata fasciata* (only AHE loci), *Rhiginia ruficoria* and *Rhiginia sp.*, and *Triatoma protracta* and *Triatoma rubrofasciata*. For each pair, we assessed the proportion of synonymous to nonsynonymous substitutions using the CODEML subprogram of *PAML* v4.9 (Yang 2007). Due to the small number of loci available for *Phymata*, only the latter two comparisons were used, but all results were provided in the supplementary data.

Based on chromosomal linkage of the loci determined by *ALiBaSeq* during dataset construction, we built a correspondence table between sex chromosome genes of the three taxa with chromosome-level assemblies: *Rhodnius*, *Triatoma*, and *Apolygus*. As the relationship between *Rhodnius* and *Triatoma* chromosomes was already established in (Mathers et al. 2020), we used the same approach of employing *BLASTP* and *MCScanX (Wang et al. 2012)* to infer chromosomal homology between *Triatoma* and *Apolygus*. BLASTP e-value cut off was set to 1e-10, match size in *MCScanX* was lowered from default 5 to 2 given the higher divergence between the species. The results were then visualized with *SynVisio* (Bandi and Gutwin 2022). To check coverage difference or lack thereof of putative autosomal and sex X loci (as determined for *Rhodnius* and *Triatoma*) in other samples, we used *BBMap* to map reads of each sample to each locus and record average read depth using the covstats parameter.

### Phylogenetic analyses

Phylogenetic analyses were carried out on the separate datasets (AHE and OMCL), as well as on the combined dataset. For each of the datasets, concatenated matrices of NT and AA data were produced. NT supermatrices were analyzed with *IQ-TREE* using standard models with partition finding for the AHE loci only (Kalyaanamoorthy et al. 2017). OMCL loci were too numerous to conduct a partition finding in a reasonable time, this similarly precluded us from trying the codon models on this data. AA supermatrices were also analyzed with *IQ-TREE*, using standard AA models and partition finding, as well as using posterior mean site frequency [PMSF (Wang et al. 2018)] analysis with LG+C20 model. Similarly to the nucleotide analyses, partition finding was not used on the OMCL loci due to computational difficulties. As PMSF results largely replicated standard AA search, we only show the latter, with the former available in supplements. Additionally, we reconstructed a final set of gene trees for all loci using *RAxML*, collapsed branches below 33% RBS. These gene trees were grouped into three datasets (AHE, OMCL, and both combined) and analyzed in *Astral* v5.6.3 (Zhang et al. 2018). Autosome-only and sex-only loci were analyzed only on nucleotide level, with reusing models for loci from the combine (AU+X) data, but for NT12 dataset with removed third codon position the model testing was performed de novo as outlined above.

### Conflict interrogation

To assess congruence and conflict among the produced topologies (Figs. 2, S1), we used the R package *Treespace (Jombart et al. 2017)* to conduct a PCoA based on RF distances calculated between trees. We used log-likelihood fit difference as described above to compute per locus phylogenetic signal, which allowed us to assess concatenation-based conflict between loci. We summed the likelihood of autosomal and sex loci to gauge the degree and direction of phylogenetic conflict between sex and non-sex-linked loci.

To investigate gene conflict beyond chromosomal linkage, we filtered out the third codon position from the alignments, computed gene trees in *RAxML* as above, and collapsed nodes with less than 50% RBS. Pairwise weighted RF distances were calculated between all gene trees and a PCoA analysis of the distances was conducted in *Treespace*. Genes were grouped into 7 clusters based on the results of the PCoA, full alignments (all codon positions) of each group were concatenated and analyzed in *IQ-TREE* to produce a concatenation-based topology for each grove.

## Acknowledgements

This research was funded by NSF DEB #1655769 “The assassin’s tale: evolutionary history of the Reduvioidea, a diverse clade of predatory and hematophagous insects” to C. Weirauch and and NSF DEB #1239788 “Phylogenomics and morphology of the hemipteroid insect orders” to K. P. Johnson. Fieldwork was partially supported by the National Geographic grant “Predators become prey: untangling spider-associated behavior and morphology within true bugs (Heteroptera)” to C. Weirauch and S. Standring. We also thank numerous colleagues for loaning or donating specimens for this project. We acknowledge anonymous reviewers and editors for providing insightful comments.

## Data availability

Raw sequence data generated for this study are available at the NCBI short-read archive, with the majority of experiments submitted under BioProject PRJNA704648. Detailed information on accession numbers is available in Table S1. Sequence assemblies, locus alignments, and other supplementary data are available from Zenodo: https://zenodo.org/record/7726314. Code used to process the data is available at GitHub: https://github.com/AlexKnyshov/reduvioid_phylogenomic_pipeline.

## Supplemental information

Supplementary Text S1. Gene content methods and results.

Supplementary Table S1. Information on samples and SRA accession numbers. Supplementary Fig S1. Individual topologies of each analysis from Fig 2. Branches are colored based on the current subfamiliar classification of the 106 ingroup taxa. Values at nodes show SH-aLRT, UFBoot, and sCF support values (IQ-TREE-based analyses) or local posterior probability (Astral-based analyses).

Supplementary Fig S2. The distribution of various locus properties in autosomal and X chromosome loci with outliers included relative to Fig. 3.

Supplementary Fig S3. IQ-TREE Binary Gene content based tree (genomes only). Supplementary Fig S4. Gains and losses of 25,954 OMCL orthogroups on pruned ML tree, partitioned into AU and X subsets. Boxes at nodes show gains and losses along internal branches. Tip gains and losses annotated on the right along with specimen sex and genome completeness proxies (% of phylogenomic loci recovered and median phylogenomic locus coverage). Since the X subset includes only 809 loci and is considerably smaller than AU, color scales are independent for each subset to show relative magnitude of change per partition.

Supplementary Fig S5. Panel (a) shows a tree inferred from aligned trimmed sequences of OBP13 (orthogroup 31851), OBP14 (orthogroup 8651), and OBP23 (orthogroup 8645) gene clusters; only OBP13 was inferred to be Triatominae-specific. Panel (b) features a UPGMA tree inferred from unaligned sequences of AGO2-like gene clusters from orthogroups 26369 (green), 4180 (purple), 2404 (red), and 3589 (turquoise), showing a Harpactorine-specific expansion. Several sequences (7 on S5a and 11 on S5b) have been deemed to be misassigned and were resigned (in color) to the appropriate OG based on more accurate phylogenetic results.

Zenodo supplements contents:

alignmentsAA.tar.gz - amino acid alignments of final loci in FASTA format

alignmentsMT.tar.gz - nucleotide alignments of mitochondrial loci in FASTA format

alignmentsNT.tar.gz - nucleotide alignments of final loci in FASTA format

alignmentsNT12.tar.gz - nucleotide alignments of final loci with excluded third codon position in FASTA format

assemblies.tar.gz - assemblies of AHE, RNA-Seq and WGS data in FASTA format

baits.tar.gz - bait sequences used to recover target loci from assemblies and compile phylogenomic alignments

bug_sex.csv - table with confirmed biological sex of specimens, used for analyses of locus coverage

coverage_data.csv - table with mean locus coverage (depth), related to Fig.3

dlnLselectNodes.csv - table with data for log-likelihood difference bar plots from Fig.2

geneContent.tar.gz - presence/absence matrices, ASR, phylogenetic inference, and other data associated with gene content analyses

geneTreeGroveAnalysis.tar.gz - data associated with the grove analysis, Fig.4

genetreesNT.tar.gz - gene trees, inferred on full final nucleotide alignments

genetreesNT12.tar.gz - gene trees, inferred on final nucleotide alignments with third codon position excluded

Gordon2017OMCLorthogroups.tar.gz - orthogroups inferred in Gordon 2017 from RNA-Seq data using OrthoMCL, in FASTA format

locus_properties.csv - table with all locus properties assessed in the present study (related to Fig.3)

selectionAnalyses.tar.gz - input data and results from the selection analyses in PAML

speciesTrees.tar.gz - main phylogenetic inference data, inputs, models, and results, related to Fig.2 and Fig.S1

syntenyAnalyses.tar.gz - input data and results from the gene synteny analyses between *Triatoma rubrofasciata* and *Apolygus lucorum*.

tar_contents.txt - list of contents of all tar files

