## Supplemental Text S1 for "Chromosome-aware phylogenomics of Assassin Bugs (Hemiptera: Reduvioidea) elucidates ancient gene conflict"

**Supplementary text S1. Gene content methods and results**

We aimed to utilize a full set of the OrthoMCL-detected homologs (N=32,675), but the curation and sequence-based analysis of a dataset this large was difficult to undertake. In addition, the OrthoMCL analysis only included transcriptomic taxa, and doing all-vs-all search for all additional to-be-included samples was unfeasible given how fragmented many of the WGS assemblies were in our dataset. After some experimentation, we settled on recoding the data as presence/absence characters and performing a phylogenetic inference on a subset of WGS samples in this matrix as well as ancestral state reconstruction for a full set of taxa included in the study.

**Methods**

**Gene content dataset construction**

As with the OMCL dataset construction, described in the main text, we started with the 32,675 orthologous groups predicted as part of Gordon (2017). To speed up the analyses while accounting for orthogroup heterogeneity, the single longest protein sequence for up to 4 taxa from each orthogroup were compiled together, and a TBLASTN search in all assemblies was performed with a cut-off e-value of 1e-10. While this approach resulted in a small fraction of misassigned sequences, it dramatically lowered missing data proportion compared to a single bait sequence search. Although we had data on transcriptomic taxa from the original OrthoMCL analysis, the transcriptomic data was reanalyzed for consistency with other samples. ALiBaSeq was used to process the search results and recover all orthologous contigs of a taxon per orthogroup.

Some orthogroups were duplicated because they contained partial sequences. This issue was flagged in initial ALiBaSeq runs, and we filtered 6202 such duplicates, resulting in a final bait-set of 26,473 orthogroups. Using these baits and 94 genomic and transcriptomic samples, ALiBaSeq recovered 25,954 loci that were used for ancestral state reconstruction analysis. Among these, 809 characters corresponded to orthogroups with genes aligned exclusively to X chromosome(s) in both triatomine reference genomes, and thus were considered tentatively X-linked characters. Preliminary analyses showed that transcriptomic taxa (both original OrthoMCL data and reanalyzed) had significantly different content from genomic samples, dramatically impacting the phylogenetic inference. Thus for the phylogenetic inference, a subset of 68 genomic taxa was chosen, limiting the possible impact of missing data on the resulting topology. Only orthogroups present in at least one taxon were retained, resulting in a smaller matrix of 23,195 loci.

We then recoded each dataset (for the inference and for the ASR) first into a count matrix (number of orthologs per taxon per OG recovered) and then into a presence/absence matrix. Since for discontiguous assemblies as well as in case of transcriptomic isoforms, it was hard to estimate the actual number of homologs correctly, we adopted the presence/absence coding as a more conservative approach. Taxa with no matches were coded as 0 while taxa with 1 or more matches coded as 1.

**Gene content phylogenetic inference**

Phylogenetic inference on the presence/absence matrix was done in IQ-TREE with appropriate model test. SH-aLRT and UFBoot support metrics (1000 resampling each) were used to assess the confidence of reconstruction.

**Gene content ancestral state reconstruction**

We also reconstructed gene content history on the NT ML tree. For that, the tree was pruned to retain gene content dataset taxa only (N=94), and a customized parsimony ancestral state reconstruction was performed on all variable characters. Given that state 0 could mean both loss of gene and missing data, equal weight parsimony was inferring too many independent gains in characters with 0 state spread throughout the tree. In gene content analysis a Dollo parsimony, which limits gene gains to a single event on a tree, was deemed more appropriate (Rogozin et al. 2006). Since we found no ready-to-use Dollo ASR solution that could handle our dataset, we opted to use asr_max_parsimony function from the R package castor, so that a custom transition matrix could be used. For each character, the initial ASR was done with equal gain/loss costs, and then cost of gain was increased by one and analysis redone until exactly one gain on the tree was inferred. A custom script then was used to transform output of asr_max_parsimony into character state changes for each branch of the tree. We separately counted gains (0>1 changes) and losses (1>0 changes) for autosomal and X-linked characters and visualized the results on the pruned NT ML tree.

**Additional analyses**

For several focal nodes, we conducted a superficial annotation of orthogroups that were gained or lost to gauge the functional aspect of gene content change by searching for homology in published Heteroptera records in the Transcriptome Shotgun Assembly Sequence Database. We further validated reconstructed gain / loss events on two gene families, odorant binding proteins (OBP) which were previously hypothesized to have Triatominae-specific expansions (Mesquita et al. 2015), and Argonaute 2 (AGO2) protein, which we detected a Harpactorinae-specific expansion. In both cases, we combined a clade-unique OG (the OG that had a clade-specific gain event) with several (3-4) OGs with the most similar sequence, and reanalyzed a combined OG data to verify orthology assessment and examine taxonomic composition of each gene tree subclade.

In the case of OBP, we added *Rhodnius prolixus* transcripts that were annotated as belonging to specific OBP clusters (Mesquita et al. 2015). Number of taxa and gene length allowed us to perform a MAFFT alignment of the sequences retrieved by ALiBaSeq, followed by trimming with trimal (-automated1 mode), and gene tree inference in RAxML under GTRGAMMA model. Seven out of 95 sequences were misassigned between two larger non-triatomine specific OGs, which we corrected in color coding using the inferred tree.

In the case of AGO2, the number of taxa (407) and divergence of rough sequences obtained via ALiBaSeq did not allow us to successfully perform a proper multiple sequence alignment. Thus, we opted to reconstruct a UPGMA tree based on pairwise distances using MAFFT (mafft --retree 0 --treeout --reorder --localpair --weighti 0 --averagelinkage). Eleven of the 407 sequences were deemed misassigned and were reassigned to the clades they were reconstructed in the UPGMA analysis (10) or removed (1).

**Results**

**Phylogenetic inference**

Model test of the presence/absence matrix yielded GTR2+FO+R5 as the best model. Phylogenetic analysis resulted in a tree surprisingly concordant with the sequence-based reconstructions (Fig. S3). This relatively simple method of analysis has also been shown to reconstruct phylogenies in cases of much deeper divergences (Pett et al. 2019). Despite a different branching order of subfamilies, monophyly of most clades from the sequence-based analyses was well supported in the gene content analysis as well (Ectrichodiinae - 100% UFBoot, Vesciinae - 100%, Peiratinae - 100%, Emesinae+Saicinae - 100%, Harpactorinae+Bactrodinae - 100%, Heteropinus+Harpactorinae+Bactrodinae - 78%, Epiroderinae+Phimophorinae - 100%, Stenopodainae - 100%, Triatominae - 100%).

**Ancestral state reconstruction**

Reconstructed gene gain and loss events are reflected in Fig. S4. Because the known X chromosome gene content is restricted to two species of Triatominae, we are not able to infer unique gene gains that may be on the X chromosome outside of this group. We also note that gene loss, especially at tips, can be due to lack of sequencing coverage and this is obvious for some taxa (e.g., most transcriptomic taxa, *Bayerus* sp., *Voconia* sp.). However, reconstruction of ortholog group gains as well as losses at deep nodes are more reliable and the total number of autosomal and X-loci retained across all taxa demonstrates that sequencing coverage was considerable even for the taxa with the lowest depth. Although X chromosome gains can not be inferred, it is notable that the ratio of autosomal losses reconstructed on nodes to X chromosome loci is almost always more frequent than would be expected if losses were randomly distributed (25,145 autosomal loci; 809 X loci; ~38 autosomal losses for every X loss). Only three of the 92 internal nodes in the phylogeny have an elevated rate of X loci compared to this ratio (*Bayerus* sp. + *Santosia* sp., *Pygolampsis* sp. + *Sastrapada* sp., and the sister clade to *Polididus* sp.) and two of those are sister pairs of male specimens where X chromosome coverage would be expected to be lower. Although this inference may be biased by the uncertain localization of ortholog groups not found in Triatominae, it may yet indicate a higher level of total conservation of X-linked loci compared to autosomal loci. Most of the gene gains occurred either at root or along the backbone of reduviids, with very few orthogroups unique to subfamily rank taxa. Some exceptions are the nodes corresponding to Peiratinae, Stenopodainae+Triatominae, and part of the higher Harpactorinae (the sister clade to *Polididus*), where each of these clades have a substantial number of gained orthogroups. This is consistent with longer branches of these clades inferred from the sequence data (Fig. 2).

We then annotated gains specific to two of the groups above, as well as Triatominae due to the importance of this group and divergence in feeding behavior which would be expected to contribute to genetic novelty.

Of 252 gained ortholog groups in Peiratinae, 143 have no significant homology to annotated proteins in Heteroptera. Others are enriched for homology to enzymes associated with salivary glands (Walker et al. 2016) like proteases (8), protease inhibitors (5) proteins with signatures for secretion (7) as well as odorant-binding proteins (3).

Of the relatively few (9) gained ortholog groups in all sampled Triatominae, two were salivary lipocalins implicated as a critical component of the anticoagulant response of these blood feeders (Santos et al. 2022) and a single protein was an odorant binding protein (OBP). As discussed in the main text, we constructed a tree of this OBP (RproOBP6) found in exclusively Triatominae species along with sequences from two other related ortholog groups that were more widespread across reduvioids, for a total of 95 sequences, excluding three reference sequences (Fig. S3a). We confirmed that all other OBPs were in fact not exclusive to Triatominae, and this one represents a unique separate ortholog group which is a promising candidate for a role in host sensing, as it has been previously determined to be expressed in the antennae of male and female insects.

Of 245 gained ortholog groups in higher Harpactorinae, 165 have no significant homology to annotated proteins in Heteroptera. Others are enriched for homology to enzymes associated with salivary glands like proteins with signatures for secretion (8), proteases (4) and “nitrophorins” (2) as well as chemosensory proteins like odorant-binding and receptor proteins (3). Interestingly, the group also appears to evolve a separate, Argonaute 2, protein involved in antiviral defense (OG1_5_26369). To confirm the accurate reconstruction of the duplication of this relatively well understood protein (Wynant et al. 2017), we created a phylogenetic tree with these sequences as well as sequences from three other ortholog groups with significant homology representing other Argonaute 2 proteins for a total of 406 sequences, excluding one reference sequence (Fig. S5b). This tree confirmed that this ortholog group (in green in Fig. S5b) was unique to the higher Harpactorinae node in question and formed a separate cluster; the other ortholog groups were widespread across Reduvioidea and included more closely related orthologs of separate genes from higher Harpactorinae in each.
