## Supplemental Figures for "Chromosome-aware phylogenomics of Assassin Bugs (Hemiptera: Reduvioidea) elucidates ancient gene conflict"

Supplementary Fig S1. Individual topologies of each analysis from Fig 2. Branches are colored based on the current subfamiliar classification of the 106 ingroup taxa. Values at nodes show SH-aLRT, UFBoot, and sCF support values (IQ-TREE-based analyses) or local posterior probability (Astral-based analyses).

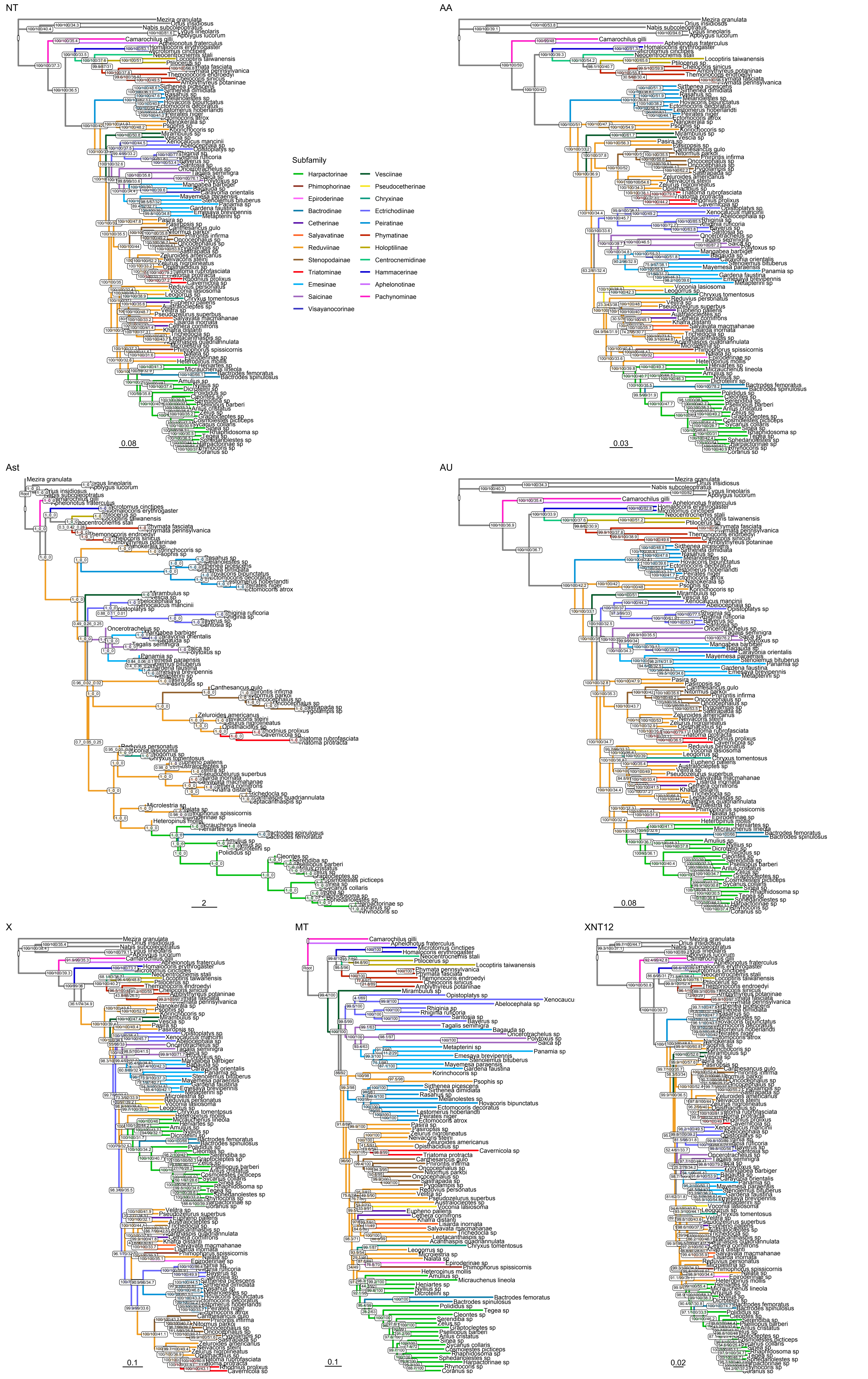
NT
AA
Ast
AU
X
MT
XNT12
Subfamily
Harpactorinae
Phimophorinae
Epiroderinae
Bactrodinae
Cetherinae
Salyavatinae
Reduviinae
Stenopodainae
Triatominae
Emesinae
Saicinae
Visayanocorinae
Vesciinae
Pseudocetherinae
Chryxinae
Ectrichodiinae
Peiratinae
Phymatinae
Holoptilinae
Centrocnemidinae
Hammacerinae
Aphelonotinae
Pachynominae
0.08
0.03
2
0.08
0.1
0.1
0.02

Supplementary Fig S2. The distribution of various locus properties in autosomal and X chromosome loci with outliers included relative to Fig. 3.

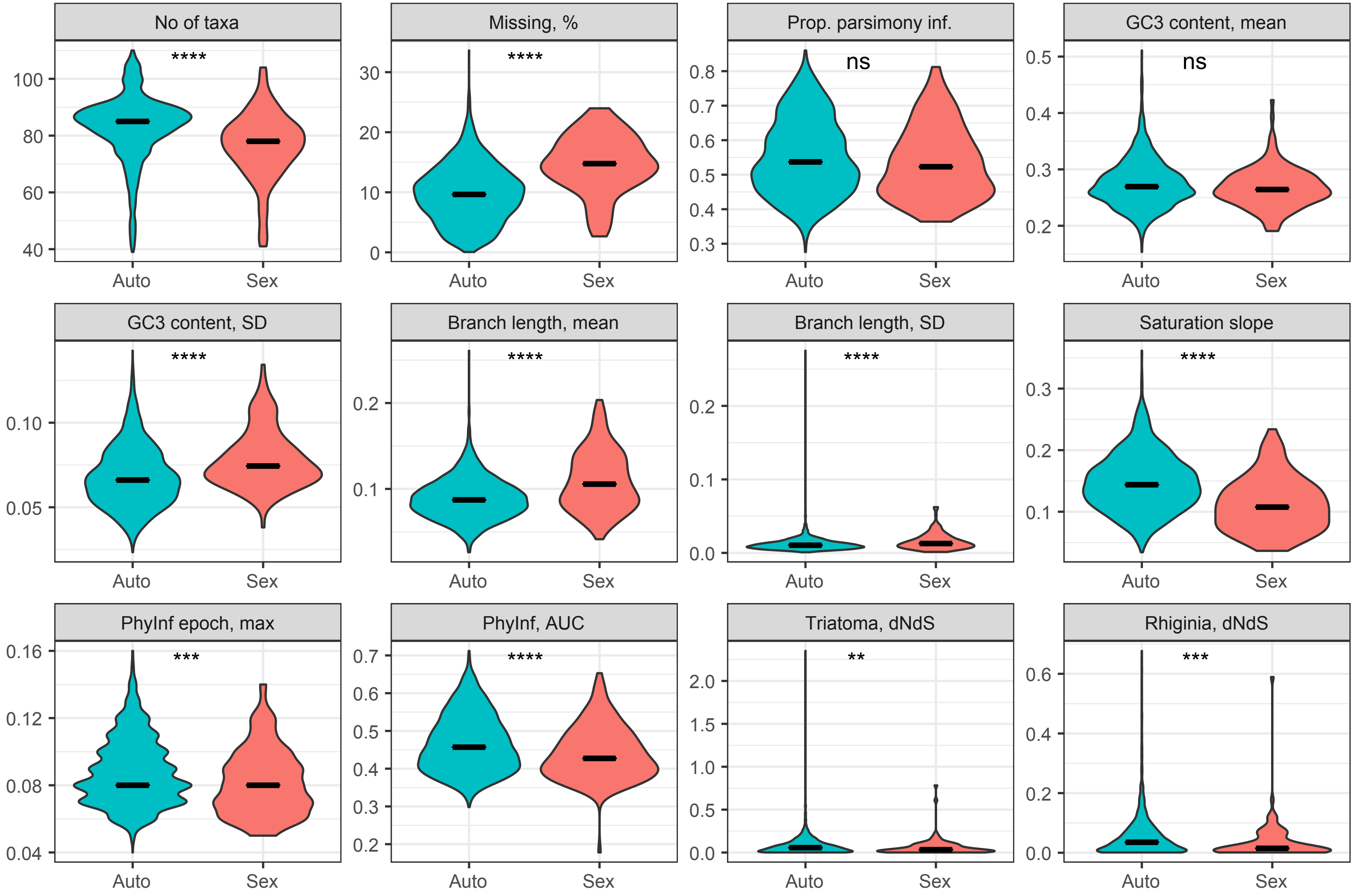

No of taxa
Missing, %
Prop. parsimony inf.
GC3 content, mean
GC3 content, SD
Branch length, mean
Branch length, SD
Saturation slope
PhyInf epoch, max
PhyInf, AUC
Triatoma, dNdS
Rhiginia, dNdS
****
ns
***
**
Auto
Sex

Supplementary Fig S3. IQ-TREE Binary Gene content based tree (genomes only).

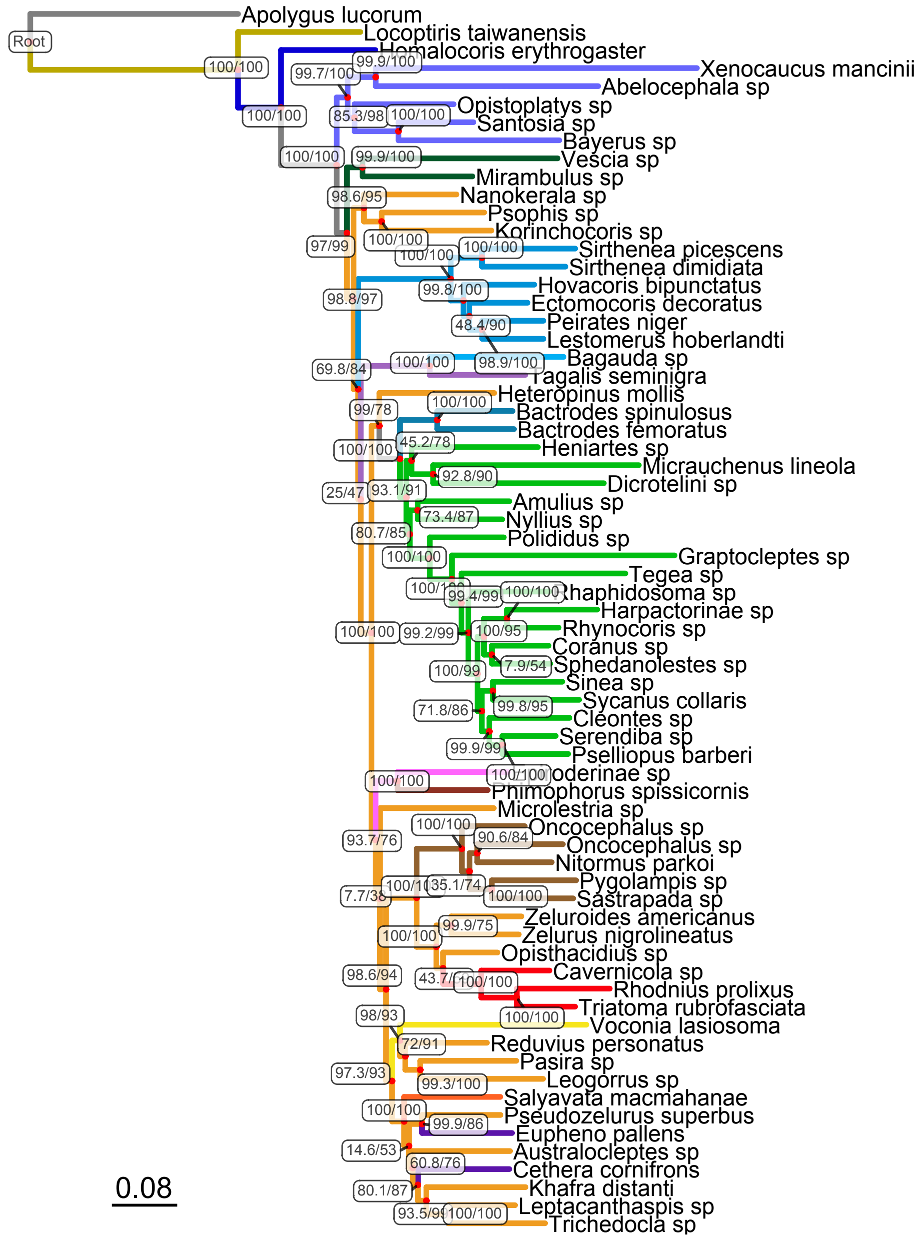
Root
Apolygus lucorum
Locoptiris taiwanensis
Homalocoris erythrogaster
Xenocaucus mancinii
Abelocephala sp
Opistoplatys sp
Santosia sp
Bayerus sp
Vescia sp
Mirambulus sp
Nanokerala sp
Psophis sp
Korinchocoris sp
Sirthenea picescens
Sirthenea dimidiata
Hovacoris bipunctatus
Ectomocoris decoratus
Peirates niger
Lestomerus hoberlandti
Bagauda sp
Tagalis seminigra
Heteropinus mollis
Bactrodes spinulosus
Bactrodes femoratus
Heniartes sp
Micrauchenus lineola
Dicrotelini sp
Amulius sp
Nyllius sp
Polididus sp
Graptocleptes sp
Tegea sp
Rhaphidosoma sp
Harpactorinae sp
Rhynocoris sp
Coranus sp
Sphedanolestes sp
Sinea sp
Sycanus collaris
Cleontes sp
Serendiba sp
Pselliopus barberi
Phimophorus spissicornis
Microlestria sp
Oncocephalus sp
Oncocephalus sp
Nitormus parkoi
Pygolampis sp
Sastrapada sp
Zeluroides americanus
Zelurus nigrolineatus
Opisthacidius sp
Cavernicola sp
Rhodnius prolixus
Triatoma rubrofasciata
Voconia lasiosoma
Reduvius personatus
Pasira sp
Leogorrus sp
Salyavata macmahanae
Pseudozelurus superbus
Eupheno pallens
Australocleptes sp
Cethera cornifrons
Khafra distanti
Leptacanthaspis sp
Trichedocla sp
100/100
100/100
99.7/100
99.9/100
85.3/98
100/100
100/100
99.9/100
98.6/95
97/99
100/100
100/100
100/100
98.8/97
99.8/100
48.4/90
100/100
98.9/100
69.8/84
99/78
100/100
100/100
45.2/78
92.8/90
25/47
93.1/91
73.4/87
80.7/85
100/100
99.4/99
100/100
100/100
99.2/99
100/95
7.9/54
100/99
99.8/95
71.8/86
99.9/99
100/100
100/100
100/100
93.7/76
90.6/84
35.1/74
7.7/38
100/100
99.9/75
100/100
98.6/94
100/100
100/100
98/93
72/91
97.3/93
99.3/100
100/100
99.9/86
14.6/53
60.8/76
80.1/87
100/100
0.08

Supplementary Fig S4. Gains and losses of 25,954 OMCL orthogroups on pruned ML tree, partitioned into AU and X subsets. Boxes at nodes show gains and losses along internal branches. Tip gains and losses annotated on the right along with specimen sex and genome completeness proxies (% of phylogenomic loci recovered and median phylogenomic locus coverage). Since the X subset includes only 809 loci and is considerably smaller than AU, color scales are independent for each subset to show relative magnitude of change per partition.

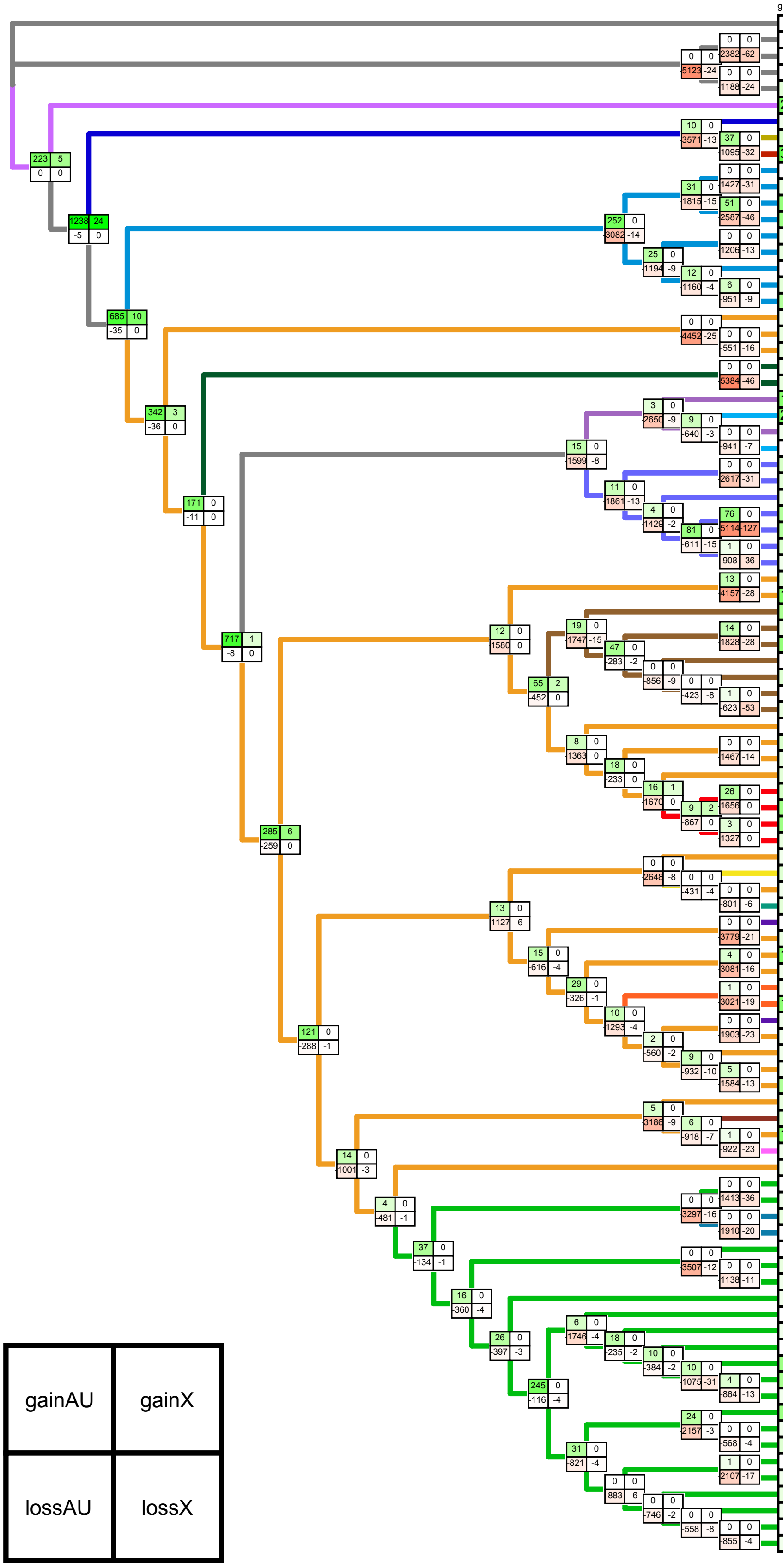


| Species | gainAU | lossAU | gainX | lossX | totalAU | totalX | sex | %Nloci | MedCov |
|---|---|---|---|---|---|---|---|---|---|
| Mezira granulata | 0 | -8898 | 0 | -163 | 8612 | 592 | | | |
| Orius insidiosus | 0 | -1474 | 0 | -32 | 8531 | 637 | | | |
| Nabis subcoleoptratus | 0 | -1444 | 0 | -72 | 8561 | 597 | | | |
| Lygus lineolaris | 0 | -2909 | 0 | -88 | 8290 | 619 | | | |
| Apolygus lucorum | 3 | -1309 | 0 | -55 | 9893 | 652 | | | |
| Aphelonotus fraterculus | 211 | -8755 | 0 | -144 | 9189 | 616 | | | |
| Homalocoris erythrogaster | 1 | -3268 | 0 | -29 | 12138 | 742 | ♀ | 84 | 9.67 |
| Locoptiris taiwanensis | 5 | -1787 | 0 | -44 | 12565 | 695 | ♂ | 78 | 15.34 |
| Phymata pennsylvanica | 396 | -2808 | 0 | -130 | 11935 | 609 | | | |
| Sirthenea picescens | 0 | -1572 | 0 | -65 | 12003 | 669 | ♂ | 88 | 7.48 |
| Sirthenea dimidiata | 2 | -1367 | 0 | -29 | 12210 | 705 | ♂ | 89 | 8.93 |
| Rasahus sp | 91 | -1624 | 0 | -66 | 10933 | 653 | | | |
| Melanolestes sp | 68 | -1994 | 0 | -34 | 10540 | 685 | | | |
| Hovacoris bipunctatus | 1 | -1045 | 0 | -27 | 13367 | 731 | ♂ | 94 | 11.35 |
| Ectomocoris decoratus | 2 | -1438 | 0 | -12 | 12975 | 746 | ♂ | 94 | 20.96 |
| Lestomerus hoberlandti | 0 | -1602 | 0 | -20 | 12867 | 747 | ♀ | 94 | 13.16 |
| Peirates niger | 3 | -777 | 0 | -20 | 12750 | 738 | ♂ | 93 | 9.65 |
| Ectomocoris atrox | 32 | -3402 | 0 | -72 | 10154 | 686 | | | |
| Nanokerala sp | 1 | -3141 | 0 | -16 | 12330 | 756 | ♂ | 94 | 7.14 |
| Psophis sp | 2 | -950 | 0 | -11 | 13971 | 745 | ♂ | 94 | 11.62 |
| Korinchocoris sp | 2 | -2338 | 0 | -78 | 12583 | 678 | ♂ | 86 | 6.14 |
| Mirambulus sp | 1 | -1890 | 0 | -70 | 12809 | 681 | ♂ | 89 | 3.58 |
| Vescia sp | 2 | -2064 | 0 | -45 | 12636 | 706 | ♀ | 71 | 4.95 |
| Oncerotrachelus sp | 151 | -6225 | 0 | -116 | 10486 | 665 | | | |
| Metapterini sp | 210 | -4295 | 0 | -127 | 11844 | 651 | | | |
| Tagalis seminigra | 1 | -1742 | 0 | -22 | 13247 | 749 | ♂ | 94 | 23.47 |
| Bagauda sp | 0 | -1900 | 0 | -21 | 13088 | 750 | | 93 | 13.23 |
| Xenocaucus mancinii | 6 | -2127 | 0 | -49 | 12619 | 697 | | 56 | 7.14 |
| Abelocephala sp | 3 | -2752 | 0 | -61 | 11991 | 685 | | 59 | 2.84 |
| Opistoplatys sp | 2 | -3605 | 0 | -38 | 12329 | 737 | | 91 | 10.65 |
| Rhiginia sp | 29 | -1729 | 0 | -65 | 8664 | 568 | | | |
| Rhiginia ruficoria | 30 | -1132 | 0 | -28 | 9262 | 605 | | | |
| Bayerus sp | 5 | -1718 | 0 | -145 | 12782 | 579 | ♂ | 76 | 2.94 |
| Santosia sp | 1 | -1725 | 0 | -18 | 12771 | 706 | ♂ | 93 | 18.09 |
| Pasira sp | 4 | -928 | 0 | -25 | 14181 | 751 | | 94 | 11.57 |
| Pasiropsis sp | 156 | -3554 | 0 | -89 | 11707 | 687 | | | |
| Canthesancus gulo | 41 | -8161 | 0 | -251 | 9014 | 540 | | | |
| Nitormus parkoi | 2 | -1018 | 0 | -43 | 14068 | 718 | ♂ | 91 | 4.66 |
| Pnirontis infirma | 99 | -3411 | 0 | -152 | 11772 | 609 | | | |
| Oncocephalus sp | 0 | -2651 | 0 | -57 | 13391 | 723 | ♀ | 85 | 4.28 |
| Oncocephalus sp | 4 | -2158 | 0 | -41 | 13465 | 731 | ♀ | 92 | 9.06 |
| Pygolampis sp | 0 | -1416 | 0 | -70 | 13581 | 649 | ♂ | 85 | 7.02 |
| Sastrapada sp | 2 | -1352 | 0 | -114 | 13647 | 605 | ♂ | 81 | 4.75 |
| Zeluroides americanus | 0 | -4186 | 0 | -22 | 13321 | 784 | | 94 | 13.28 |
| Neivacoris steini | 5 | -1778 | 0 | -148 | 14052 | 644 | | 74 | 2.75 |
| Zelurus nigrolineatus | 2 | -2082 | 0 | -15 | 13745 | 777 | ♂ | 94 | 6.48 |
| Opisthacidius sp | 3 | -2044 | 0 | -12 | 13597 | 795 | ♀ | 96 | 13.67 |
| Triatoma rubrofasciata | 7 | -1403 | 0 | 0 | 11754 | 809 | | | |
| Triatoma protracta | 62 | -2125 | 0 | -82 | 11087 | 727 | | | |
| Rhodnius prolixus | 71 | -2083 | 0 | 0 | 11444 | 809 | | | |
| Cavernicola sp | 2 | -987 | 0 | -17 | 12471 | 792 | ♀ | 94 | 11.83 |
| Reduvius personatus | 1 | -3484 | 0 | -19 | 13405 | 770 | ♀ | 95 | 10.12 |
| Voconia lasiosoma | 4 | -3256 | 0 | -173 | 13205 | 612 | ♂ | 73 | 2.56 |
| Leogorrus sp | 1 | -1533 | 0 | -28 | 14124 | 751 | ♀ | 88 | 5.75 |
| Chryxus tomentosus | 3 | -2290 | 0 | -43 | 13369 | 736 | ♀ | 87 | 3.92 |
| Eupheno pallens | 0 | -2044 | 0 | -15 | 13112 | 757 | ♀ | 94 | 13.99 |
| Australocleptes sp | 2 | -1172 | 0 | -25 | 13986 | 747 | ♂ | 94 | 10.33 |
| Velitra sp | 153 | -4176 | 0 | -124 | 11538 | 652 | | | |
| Pseudozelurus superbus | 1 | -2111 | 0 | -28 | 13451 | 748 | ♂ | 94 | 1.85 |
| Salyavata macmahanae | 1 | -1446 | 0 | -25 | 12890 | 744 | ♀ | 94 | 8.11 |
| Lisarda inornata | 119 | -3866 | 0 | -128 | 10588 | 641 | | | |
| Cethera cornifrons | 1 | -1747 | 0 | -23 | 13148 | 740 | | 92 | 4.9 |
| Khafra distanti | 1 | -1227 | 0 | -99 | 13668 | 664 | | 88 | 3.67 |
| Trichedocla sp | 1 | -1380 | 0 | -50 | 14495 | 726 | | 93 | 6.34 |
| Leptacanthaspis sp | 1 | -784 | 0 | -31 | 13512 | 732 | | 94 | 8.29 |
| Acanthaspis quadriannulata | 36 | -5020 | 0 | -154 | 9311 | 609 | | | |
| Microlestria sp | 2 | -2646 | 0 | -31 | 13838 | 760 | | 95 | 11.08 |
| Phimophorus spissicornis | 5 | -2661 | 0 | -22 | 12914 | 762 | | 94 | 5.37 |
| Nalata sp | 139 | -3291 | 0 | -116 | 11497 | 645 | | | |
| Epiroderinae sp | 1 | -1390 | 0 | -33 | 13260 | 728 | ♂ | 93 | 8.36 |
| Heteropinus mollis | 1 | -5740 | 0 | -62 | 13447 | 737 | ♂ | 88 | 4.79 |
| Heniartes sp | 0 | -1432 | 0 | -22 | 12947 | 724 | | 84 | 8.64 |
| Micrauchenus lineola | 1 | -2595 | 0 | -174 | 11785 | 572 | | 67 | 6.36 |
| Bactrodes femoratus | 2 | -1316 | 0 | -20 | 12568 | 742 | | 93 | 28.92 |
| Bactrodes spinulosus | 0 | -1400 | 0 | -19 | 12482 | 743 | | 92 | 27.94 |
| Amulius sp | 0 | -2080 | 0 | -10 | 13158 | 772 | | 94 | 24.22 |
| Nyllius sp | 2 | -1529 | 0 | -17 | 12573 | 754 | | 94 | 6.14 |
| Dicrotelini sp | 1 | -2120 | 0 | -66 | 11981 | 705 | ♀ | 69 | 5.92 |
| Polididus sp | 1 | -5340 | 0 | -48 | 13035 | 743 | ♀ | 93 | 7.08 |
| Cleontes sp | 0 | -3812 | 0 | -32 | 12951 | 751 | | 95 | 11.03 |
| Serendiba sp | 1 | -2873 | 0 | -30 | 13674 | 751 | ♂ | 96 | 12.24 |
| Pselliopus barberi | 0 | -2087 | 0 | -19 | 14085 | 760 | ♂ | 95 | 8.75 |
| Arilus cristatus | 70 | -4242 | 0 | -147 | 10935 | 601 | | | |
| Zelus sp | 48 | -3890 | 0 | -60 | 10405 | 675 | | | |
| Graptocleptes sp | 1 | -1565 | 0 | -163 | 12683 | 572 | | 67 | 3.65 |
| Cosmolestes picticeps | 75 | -4375 | 0 | -143 | 11280 | 637 | | | |
| Sycanus collaris | 1 | -1603 | 0 | -62 | 13410 | 714 | | 88 | 10.61 |
| Sinea sp | 4 | -1223 | 0 | -13 | 13793 | 763 | ♀ | 95 | 12.94 |
| Rhaphidosoma sp | 0 | -1130 | 0 | -13 | 13594 | 747 | ♂ | 93 | 11.34 |
| Tegea sp | 1 | -2444 | 0 | -93 | 12281 | 667 | ♂ | 82 | 9.05 |
| Sphedanolestes sp | 2 | -2478 | 0 | -24 | 13608 | 751 | ♀ | 95 | 10.7 |
| Harpactorinae sp | 1 | -1938 | 0 | -36 | 13589 | 731 | ♂ | 93 | 10.51 |
| Rhynocoris sp | 0 | -917 | 0 | -10 | 13754 | 753 | ♀ | 94 | 15.3 |
| Coranus sp | 1 | -1457 | 0 | -57 | 13215 | 706 | ♂ | 93 | 8.67 |

Supplementary Fig S5. Panel (a) shows a tree inferred from aligned trimmed sequences of OBP13 (orthogroup 31851), OBP14 (orthogroup 8651), and OBP23 (orthogroup 8645) gene clusters; only OBP13 was inferred to be Triatominae-specific. Panel (b) features a UPGMA tree inferred from unaligned sequences of AGO2-like gene clusters from orthogroups 26369, 4180, 2404, and 3589, showing a Harpactorine-specific expansion.

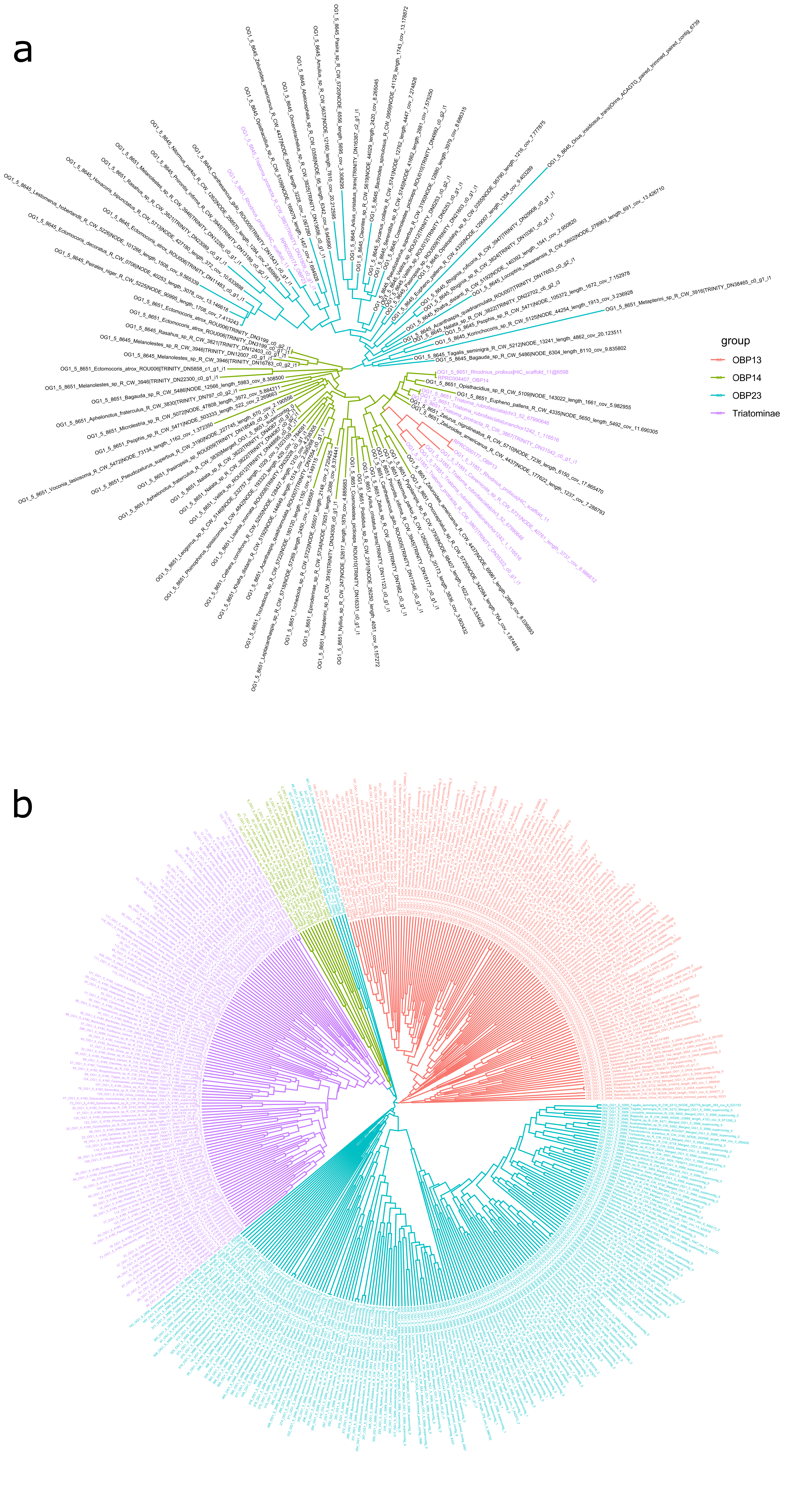
a
group
OBP13
OBP14
OBP23
Triatominae
b
